# Energy extraction from air: structural basis of atmospheric hydrogen oxidation

**DOI:** 10.1101/2022.10.09.511488

**Authors:** Rhys Grinter, Ashleigh Kropp, Hari Venugopal, Moritz Senger, Jack Badley, Princess Cabotaje, Sven T. Stripp, Christopher K. Barlow, Matthew Belousoff, Gregory M. Cook, Kylie A. Vincent, Ralf B. Schittenhelm, Syma Khalid, Gustav Berggren, Chris Greening

**Affiliations:** Department of Microbiology, Biomedicine Discovery Institute, Monash University, Clayton, VIC 3800, Australia; Ramaciotti Centre for Cryo-Electron Microscopy, Monash University, Clayton, VIC 3168, Australia; Department of Chemistry, Ångström Laboratory, Uppsala University, Box 523, 75120 Uppsala; Department of Biochemistry, University of Oxford, Oxford, OX1 3QU, UK; Department of Physics, Experimental Molecular Biophysics, Freie Universität Berlin, 14195 Berlin, Germany; Department of Biochemistry, Monash Biomedicine Discovery Institute, Monash University, Clayton, VIC 3800, Australia; Monash Proteomics & Metabolomics Facility, Monash Biomedicine Discovery Institute, Monash University, Clayton, VIC 3800, Australia; Centre for Electron Microscopy of Membrane Proteins, Monash Institute of Pharmaceutical Sciences, Parkville, 3052, Victoria, Australia; Department of Microbiology and Immunology, University of Otago, Dunedin 9016, New Zealand; Inorganic Chemistry Laboratory, Department of Chemistry, University of Oxford, Oxford, OX1 3QR, United Kingdom; Securing Antarctica’s Environmental Future, Monash University, Clayton, VIC 3800; Centre to Impact AMR, Monash University, Clayton, VIC 3800

**Author notes:** These authors contributed equally to this work (Rhys Grinter, Ashleigh Kropp).

## Abstract

Diverse aerobic bacteria use atmospheric H_2_ as an energy source for growth and survival. This recently discovered yet globally significant process regulates the composition of the atmosphere, enhances soil biodiversity, and drives primary production in certain extreme environments. Atmospheric H_2_ oxidation has been attributed to still uncharacterised members of the [NiFe]-hydrogenase superfamily. However, it is unresolved how these enzymes overcome the extraordinary catalytic challenge of selectively oxidizing picomolar levels of H_2_ amid ambient levels of the catalytic poison O_2_, and how the derived electrons are transferred to the respiratory chain. Here we determined the 1.52 Å resolution CryoEM structure of the mycobacterial hydrogenase Huc and investigated its mechanism by integrating kinetics, electrochemistry, spectroscopy, mass spectrometry, and molecular dynamics simulations. Purified Huc is an oxygen-insensitive enzyme that couples the oxidation of atmospheric H_2_ at its large subunit to the hydrogenation of the respiratory electron carrier menaquinone at its small subunit. The enzyme uses a narrow hydrophobic gas channel to selectively bind atmospheric H_2_ at the expense of O_2_, while three [3Fe-4S] clusters and their unusual ligation by a D-histidine modulate the electrochemical properties of the enzyme such that atmospheric H_2_ oxidation is energetically feasible. Huc forms an 833 kDa complex composed of an octamer of catalytic subunits around a membrane-associated central stalk, which extracts and transports menaquinone a remarkable 94 Å from the membrane, enabling its reduction. These findings provide a mechanistic basis for the biogeochemically and ecologically critical process of atmospheric H_2_ oxidation. Through the first characterisation of a group 2 [NiFe]-hydrogenase, we also uncover a novel mode of energy coupling dependent on long-range quinone transport and pave way for the development of biocatalysts that oxidize H_2_ in ambient air.

## Introduction

The oxidation of atmospheric hydrogen (H_2_) by soils is a key biogeochemical process that shapes the redox state of the atmosphere [1, 2]. Thought until recently to be an abiotic process, it is now recognised that diverse aerobic bacteria from at least nine phyla oxidise atmospheric H_2_ and together account for 75% (~60 teragrams) of the total H_2_ removed from the atmosphere annually [3–7]. Atmospheric H_2_ oxidation provides bacteria with a supplemental energy source in nutrient-limited soil environments, allowing them to either grow mixotrophically [6, 8–11], or persist on air alone in a dormant but viable state for long periods [4, 5, 12–18]. For example, both *Mycobacterium* cells and *Streptomyces* spores survive during starvation by transferring electrons through an aerobic respiratory chain from atmospheric H_2_ to O_2_ [6, 14, 19, 20]. While the ability to oxidize atmospheric H_2_ is widespread in bacteria from diverse environments [16], some ecosystems such as hyper-arid polar soils appear to be primarily driven by atmospheric energy sources [7, 15, 16, 21].

The oxidation of atmospheric H_2_ is a remarkable challenge that no current chemical catalysts can achieve: it requires the selective oxidation of low concentrations of substrate (530 ppbv) present in an atmosphere rich with the catalytic poison O_2_ (~21%) [22, 23]. Group 1 [NiFe]-hydrogenases are a family of membrane-bound H_2_-oxidizing metalloenzymes that support aerobic and anaerobic growth of bacteria in H_2_-rich environments; however, these enzymes are incapable of atmospheric H_2_ oxidation given they have a low affinity for H_2_ (K_m_ > 500 nM) and are either reversibly or irreversibly inhibited by O_2_ [24–28]. More recently, several high-affinity lineages of group 1 and 2 [NiFe]-hydrogenases have been identified that input atmospheric H_2_-derived electrons into the aerobic respiratory chain [4, 5, 21, 29, 30]. Whole-cell studies suggest these enzymes have a significantly higher apparent affinity for H_2_ (K_m_ 30 to 200 nM) and appear to be insensitive to inhibition by O_2_ [4, 5, 10, 12, 13, 31, 32]. However, given these hydrogenases have yet to be isolated, it remains unknown how they have evolved to selectively oxidize H_2_, tolerate exposure to O_2_, and interact with the electron transport chain. Notably, it is debated whether the hydrogenases responsible for atmospheric H_2_ have an inherently high affinity or whether their affinity is modulated by their interactions with the respiratory chain [7, 27].

To address these knowledge gaps, we investigated the structural and mechanistic basis of atmospheric H_2_ oxidation in the aerobic bacterium *Mycobacterium smegmatis.* This bacterium possesses two phylogenetically distinct hydrogenases, designated Huc (group 2a) and Hhy (group 1h), that both oxidize H_2_ to subatmospheric levels [5, 20, 32]. We isolated Huc directly from *M. smegmatis* and determined the structural and biochemical basis for its oxidation of atmospheric H_2_.

### Mycobacterial Huc is a high-affinity, oxygen-insensitive hydrogenase that specifically reduces menaquinone analogues

To natively express and purify Huc, we introduced a Strep-tag II at the N-terminus of *hucS* (the gene encoding the Huc small subunit) into the chromosome of a Huc overexpressing strain of *M. smegmatis* [33]. Huc was isolated from cell lysates by StrepTactin affinity chromatography and size exclusion chromatography. When analysed by SDS-PAGE, Huc consisted of three protein subunits corresponding to HucL (~58 kDa), HucS-2×Strep (~39 kDa), and a third unknown subunit (~18 kDa) (Fig. S1a). The band corresponding to the third subunit was excised and identified by tandem mass spectrometry as the uncharacterised protein MSMEG_2261, encoded by an open reading frame in the *huc* operon directly upstream of *hucS* (Fig. S1b) [20]. The co-purification of MSMEG_2261 with the hydrogenase large and small subunits suggests it is an additional component of the Huc complex, which we designate HucM. As previously observed for Huc in *M. smegmatis* cell lysates, on a native gel, the purified Huc complex migrates at a molecular mass of ~800-900 kDa (Fig. S1c) [32]. Huc is red-brown, consistent with the presence of multiple metal clusters, and is highly stable at room temperature with a melting temperature of 78.3°C (Fig. S1d). To investigate the role of HucM in the Huc complex, we deleted the *hucM* gene from an *M. smegmatis* strain in which Hhy, the other high-affinity hydrogenase encoded by this bacterium, is inactive. Activity staining of cell lysates shows that Huc is still active in this mutant, but the presence of a high molecular weight species was abolished, suggesting HucM plays an important role in the formation of the Huc oligomer (Fig. S1e).

While Huc can oxidise atmospheric H_2_ in whole cells, it was unclear whether this enzyme has an inherently high substrate affinity or if this property results from coupling to the respiratory electron transport chain [5]. To resolve this question, we tested the ability of purified Huc to oxidise 10 ppm H_2_ supplemented to an ambient air headspace. Huc rapidly consumed H_2_ to below atmospheric concentrations with nitroblue tetrazolium chloride (NBT) as the electron acceptor (Fig. 1a). Kinetic analysis of purified Huc indicates that the enzyme is adapted to oxidize atmospheric H_2_, with a high affinity (K_m_ = 129 nM) and low H_2_ threshold (280 pM), but slow turnover (K_cat_ = 7.05 s^−1^) (Fig.1b & 1c, Table S1). This is the first report of atmospheric H_2_ oxidation and high-affinity substrate kinetics by a purified hydrogenase. The kinetics of the purified hydrogenase are remarkably similar to that of *M. smegmatis* mutants harbouring Huc as the sole hydrogenase (K_m_ = 184 nM, threshold = 133 pM), suggesting the hydrogenase itself is the primary determinant of whole-cell substrate affinity [5]. The ability of Huc to oxidize H_2_ to this concentration in ambient air also indicates high tolerance to inhibition by O_2_. To assess the extent of Huc O_2_ tolerance, we amperometrically measured its rate of H_2_ oxidation in buffer containing O_2_ at 0%, 10% (~0.12 mM), and 100% (~1.2 mM) saturation [34]. No significant difference in rate or affinity of Huc mediated H_2_ oxidation was observed with varying O_2_ concentrations, indicating Huc is insensitive to inhibition by this gas (Fig.1b). Nevertheless, Huc does transfer electrons directly to O_2_ (Fig. 1c). Similar observations have been made for O_2_-tolerant enzymes, i.e. enzymes that are reversibly inhibited by O_2_, such as Hyd-1 of *E. coli* [35]; in these enzymes, O_2_ binds at the active site and is reduced to innocuous water through the reverse transfer of four electrons [25, 26]. Huc potentially functions by a similar mechanism, for example, as a functional dimer in which H_2_-derived electrons are transferred from an active hydrogenase to an inhibited O_2_-bound partner [36], though it’s alternatively possible O_2_ binds and is reduced at the electron acceptor site or another site. Irrespectively, O_2_ does not inhibit H_2_ oxidation across a range of concentrations in contrast to O_2_-tolerant enzymes and hence Huc can be classified as O_2_-insensitive enzyme.

**Figure 1.**
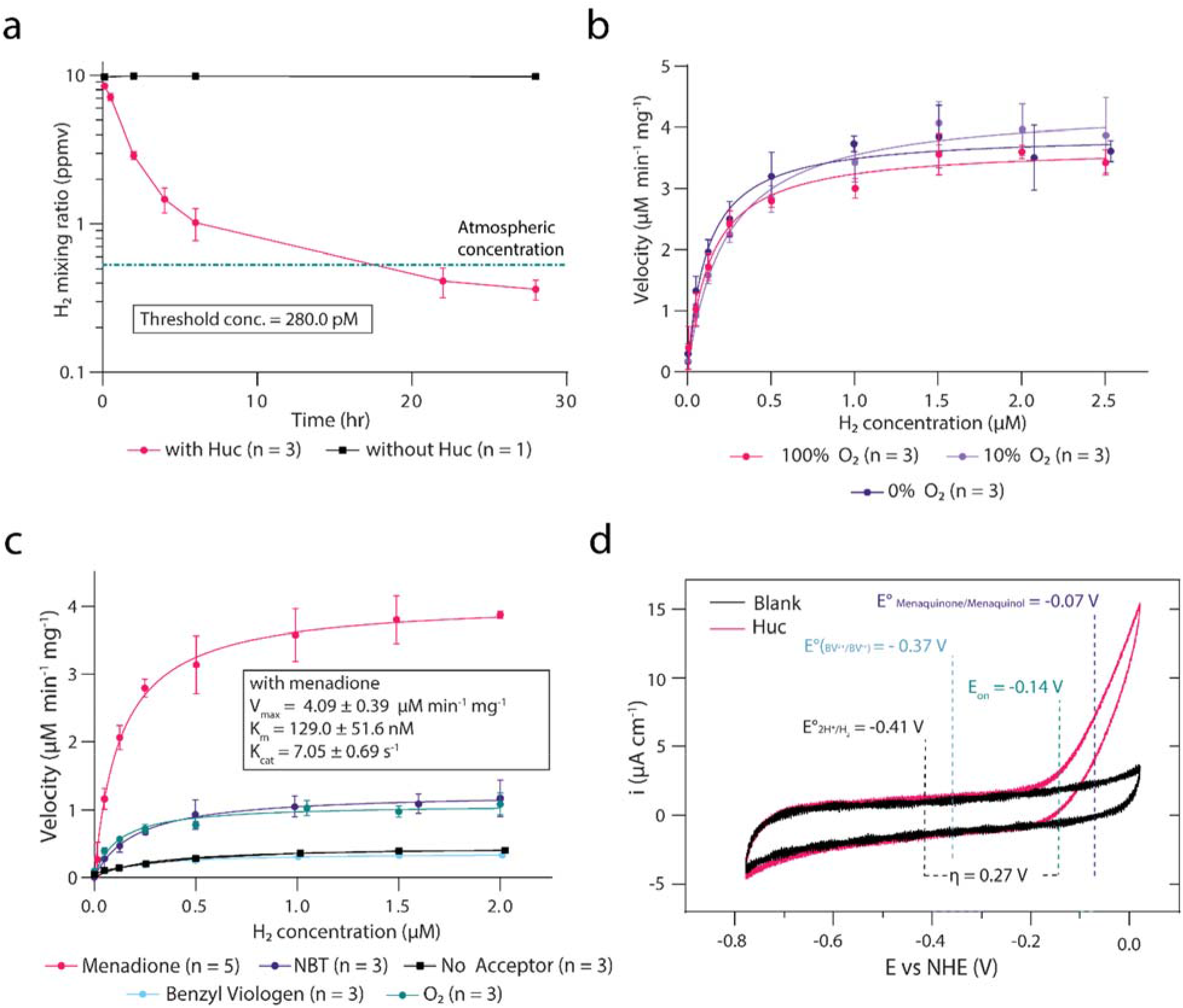
Huc is a high affinity O_2_ insensitive hydrogenase. (a) Gas chromatography analysis of the H_2_ concentration of the headspace of sealed vials containing Huc or buffer control (without Huc), showing Huc can oxidize H_2_ to below atmospheric concentrations (green line). (b) Michaelis–Menten kinetics of Huc H_2_ oxidation in buffer with different percentages of O_2_ saturation, amended with H_2_ at 1% saturation (~8 μM @ 25°C), with 200 μM menadione as electron acceptor (0% O_2_ = degassed buffer with traces of O_2_). (c) Michaelis–Menten kinetics of Huc H_2_ consumption in buffer with different electron acceptors, with 1% H_2_ saturation (~8 μM @ 25°C), degassed but with traces of O_2_. (d) Cyclic voltammogram of an immobilized Huc protein film in a 100% H_2_ atmosphere. Eon indicates the Huc onset potential, determined from the average of four experiments (a single representative curve is shown for clarity). η shows the estimated Huc overpotential compared to the hydrogen redox couple potential (E°H^+^/H_2_). The midpoint potentials of benzyl viologen (BV) and menaquinone (MQ) are indicated.

In *M. smegmatis* cells, electrons from H_2_ oxidation by Huc enter the respiratory electron transport chain via the reduction of menaquinone, the major respiratory quinone of mycobacteria [32]. However, respiratory quinones are hydrophobic and are generally reduced within the membrane, via specialized membrane proteins [37]. Whether Huc reduces menaquinone directly or via an unknown membrane protein intermediate is unknown [32]. To test if menaquinone is the immediate electron acceptor of Huc, we determined its enzyme kinetics with either menadione (representing the redox headgroup of menaquinone), or the redox-active dyes NBT or benzyl viologen (BV) as electron acceptors. Consistent with menaquinone acting as its physiological electron acceptor, Huc displayed a much higher maximum velocity with menadione (E° −74 mV) as its electron acceptor than NBT (E° −80 mV), despite their comparable midpoint potentials (Fig. 1c, Table S1) [38, 39]. The low potential electron acceptor BV (E° −374 mV) did not induce H_2_ oxidation compared to the buffer control, indicating it does not accept electrons from Huc (Fig. 1c) [40]. This is consistent with the role of Huc in oxidizing H_2_ at low concentrations, as the higher redox potential of the 2H^+^/H_2_ redox couple under ambient conditions (E° −136 mV) would preclude the reduction of low potential substrates, and thus Huc is likely tuned to higher redox potentials. Consistently, protein film electrochemistry (PFE) showed that Huc has an onset potential for H_2_ oxidation of about −140 mV (vs NHE). This is 270 mV more positive than the thermodynamic potential of the 2H^+^/H_2_ couple under the conditions tested and the highest overpotential for an [NiFe]- hydrogenase recorded to date (Fig. 1d) [35, 41–44]. Huc is an irreversible H_2_-oxidising enzyme and did not display any current attributable to H_2_ production under negative potentials during PFE experiments (Fig. 1d).

### The large and small subunits of Huc oligomerise around a central membrane-associated stalk

To determine the structure of Huc, we performed CryoEM imaging and single-particle reconstruction of the purified 833 kDa complex. CryoEM micrographs and resulting class averages revealed a four-leaf clover-shaped molecule, which in some cases is associated with membrane vesicles via a stalk-like protrusion (Fig. S2a,b). In the Huc-overexpressing strain used for purification, Huc activity is mainly associated with the soluble fraction, yet previous reports indicate that Huc predominantly associates with the cellular membrane in wildtype *M. smegmatis* [5, 32], which is consistent with the presence of membrane-associated Huc in our sample. 3D reconstruction of the entire Huc complex yielded maps at a 2.19 Å-overall resolution (Fig. S2c,d,e, Table S2), showing that each of the four lobes of the Huc molecule is composed of two Huc protomers each consisting of a HucS and HucL subunit (Fig. 2a). The four Huc lobes are bound to a scaffold formed by four molecules of HucM, an elongated α-helical protein, which intertwines to form a cage-like structure (Fig. 2b, Fig S3a,b). Because of flexibility exhibited by this complex (Fig S2f, Mov. S1), we performed 3D reconstruction of the individual Huc lobe from two datasets, obtaining resolutions of 1.52 Å and 1.67 Å (Fig S3c,d, Fig S4, Fig S5, Table S2) and revealing the asymmetric unit of the C4 symmetrical Huc oligomer (Fig. 2c). This is the highest resolution CryoEM protein structure reported to date other than ferritin [45]. The overall fold of the HucSL heterodimer is similar to that of other structurally characterized hydrogenases, with an all-residue backbone RMSD of 3.7 Å and 2.8 Å to hydrogenases from *Desulfovibrio vulgaris* Miyazaki F (PDB ID = 4U9H) and *Cupriavidus necator* H16 (PDB ID = 5AA5) respectively [27, 28]. Consistently, the architecture of the [NiFe] active site is the same as previously reported (Fig. 3a) [46]. All three iron-sulfur clusters of the small subunit have a [3Fe-4S] configuration (Fig. 3b), in contrast to other characterized hydrogenases, including O_2_-tolerant variants, that possess one or more [4Fe-4S] clusters [25, 46]. [3Fe-4S] clusters generally undergo their +/0 redox transition at a higher potential than [4Fe-4S] clusters [47], meaning these clusters likely play a major role in tuning Huc to its high overpotential [48].

**Figure 2.**
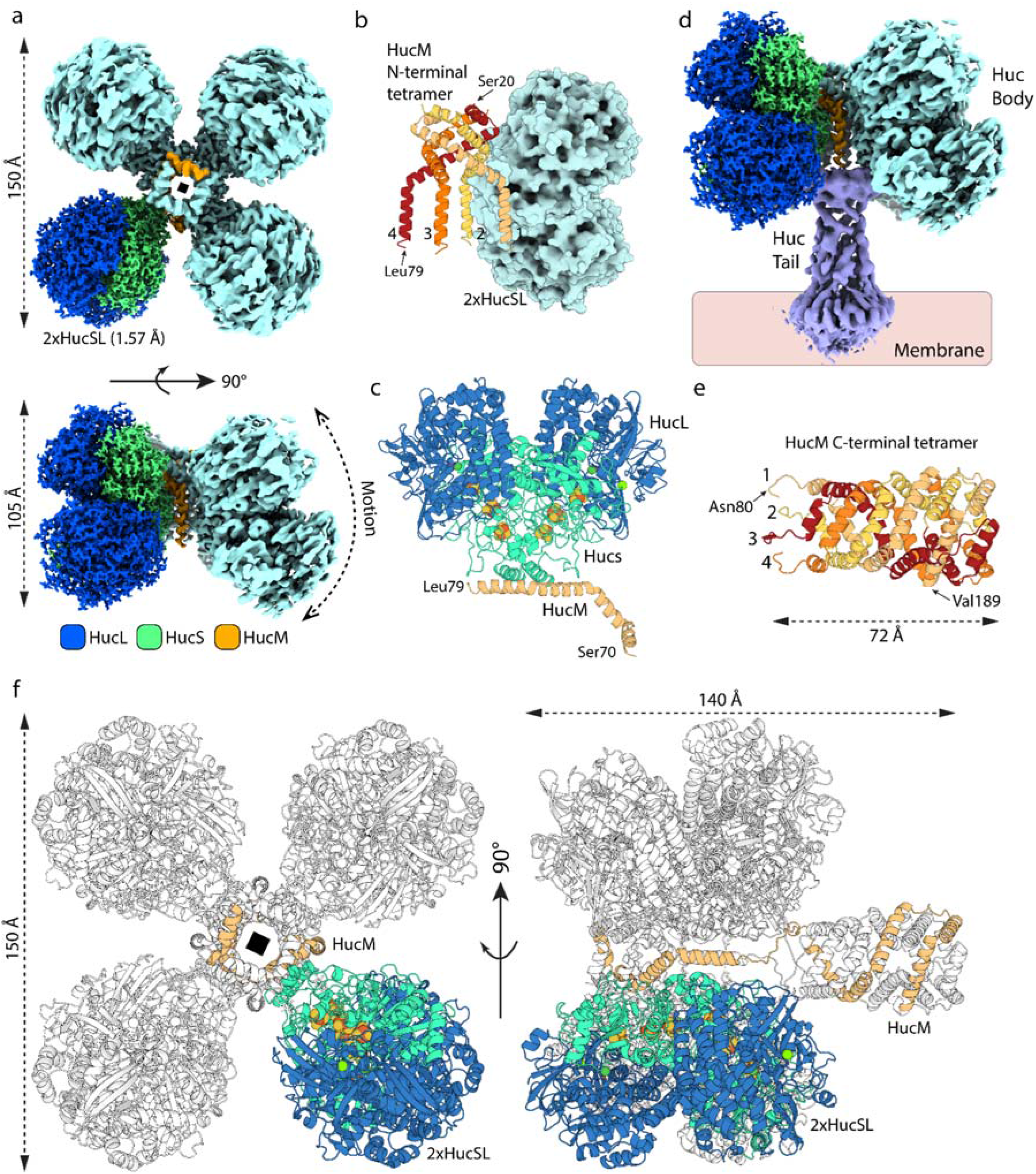
Huc forms an 833 kDa oligomer composed of three subunits HucS, HucL, and HucM. (a) CryoEM density maps of the Huc oligomer showing its four lobes that each contain two HucSL dimers, with four centrally located HucM subunits holding the oligomer together. On one lobe, HucL and HucS subunits are coloured blue and green respectively, and one HucM molecule is coloured orange. (b) A cartoon representation of the central tetramer formed by four HucM subunits, with a single HucS_2_L_2_ lobe shown as a surface model for context. (c) A cartoon representation of the asymmetric unit of the Huc oligomer. (d) A composite CryoEM density map, showing the Huc ‘body’ region as in panel a, and the ‘stalk’ region and associated lipid bilayer map reconstructed separately. (e) A cartoon representation of the AlphaFold2 model of the C-terminal region of HucM (residues 80 to 189), showing that it forms a tetrameric coiled-coil tube. (f) A cartoon representation showing the full Huc complex reconstructed from the CryoEM structure and Alphafold HucM C-terminal model fitted to the stalk region CryoEM map.

**Figure 3.**
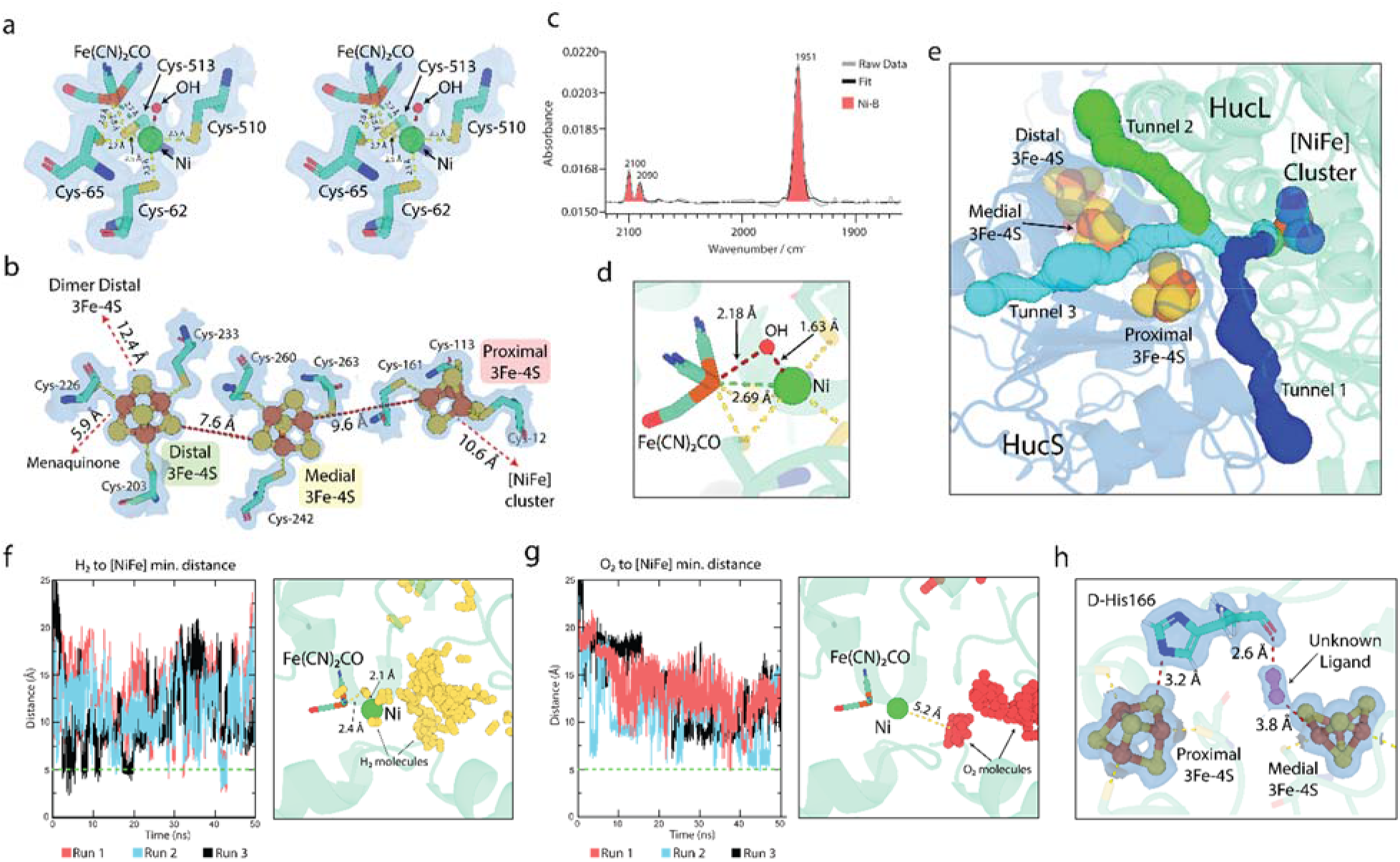
The Huc catalytic cluster adopts a non-inhibitory Ni-B state under ambient conditions. (a) A stereoview of the coordination structure of the Huc [NiFe] cluster under ambient air. All coordinating cysteines originate from the HucL subunit (b) The coordination structure and distances between the three HucS [3Fe-4S] clusters. All coordinating cysteines originate from the HucS subunit. The corresponding electron density map of the regions shown in panels a and b is shown as a transparent blue surface at 5 σ. (c) FTIR spectra of Huc isolated in ambient air and collected under N_2_, showing that the [NiFe] active site of the enzyme adopts the Ni-B state. (d) The Huc [NiFe] active site showing the relative distances between the Ni-B state hydroxyl ligand and the Ni and Fe ions of the cluster. (e) A zoomed view of the HucSL dimer showing the location and width of the hydrophobic gas channels that provide substrate access to the [NiFe] active site. See Table S3 for tunnel width statistics. (f) A distance-over-time plot showing the closest distance achieved by H_2_ to the Huc [NiFe] active site during molecular dynamics simulations of HucSL in the presence of H_2_ (left panel), the location of H_2_ molecules in a representative subset (~100 frames from Runs 1, 2, and 3) of simulation frames encompassing regions when H_2_ molecules are in the closet proximity to the active site (right panel). (g) Distance-over-time and representative closest O_2_ locations for simulations of HucSL performed in the presence of O_2_, as in panel f. (h) The location of D-histidine 166 from HucL relative to the proximal and medial [3Fe-4S] clusters, and the location of the unknown ligand. Electron density maps from the high-resolution reconstruction and shown as a transparent blue surface at 5 σ.

Out of the 188 amino acids of HucM, only residues 20-84 could be modelled into CryoEM density, with diffuse electron density corresponding to the remaining C-terminal region of HucM forming the aforementioned stalk-like protrusion. To better resolve this region of the structure, we masked it and performed signal subtraction on the remainder of the Huc structure followed by focused refinement and 3D classification (Fig S6, Table S2). This processing showed that the HucM C-terminal region forms a tube connecting the main body of the Huc oligomer to the membrane (Fig. 2d, Fig. S7a,c). The electron density indicates that this tube connects to the body of Huc through the coiling of the four HucM molecules, with this structure enclosing an internal chamber at the centre of Huc (Fig. S7b,c). We used AlphaFold to model the structure of the HucM C-terminal region [49]. Consistent with the CryoEM data, the resulting tetrameric assembly forms a tube composed of a symmetrical coiled-coil of α-helices composed of the four HucM molecules (Fig. 2e). These α-helices are amphipathic, with the hydrophobic face pointing inwards, lining the tube entirely with hydrophobic residues (Fig. S7d). Interaction of the tube with the membrane is facilitated by an outward-facing helix at the tube terminus containing arginine and lysine residues that facilitate interaction with the phospholipid head groups, and a tryptophan residue that may stabilize the tube at the membrane surface through partial insertion into the bilayer (Fig. S7e). By docking the model of the HucM tetramer into the map for the Huc stalk, we resolved the structure of the complete Huc complex (Fig. 2f, Mov. S2).

### Narrow hydrophobic gas channels and high-potential iron-sulfur clusters underlie high-affinity kinetics and O_2_ insensitivity

Characterization of purified Huc demonstrates that it is inherently capable of H_2_ oxidation at sub-atmospheric concentrations and is insensitive to inhibition by O_2_. These properties are remarkable for a hydrogenase and are highly desirable for the development of biocatalysts that oxidize H_2_ under aerobic conditions [50–52]. In previously studied respiratory O_2_-tolerant hydrogenases, it is proposed that O_2_ binds to the [NiFe] active site and is rapidly reduced to water by four electrons derived from the three iron-sulfur clusters of the small subunit [25, 26, 53, 54]. Three of these electrons are provided by ordinary redox transitions of the reduced clusters that reduce the bound O_2_ to hydroxide, thus preventing the formation of reactive oxygen species, and a fourth electron originating from an additional high potential redox transition by the unusual [4Fe-3S] proximal cluster further reduces the hydroxyl to water removing it from the [NiFe] cluster [25, 26, 55]. However, as Huc exclusively binds [3Fe-4S] clusters it is unable to fully reduce O_2_ in this way and must resist O_2_ inactivation by a distinct mechanism. As with several other [NiFe]-hydrogenases, the distal Fe-S clusters of the HucS_2_L_2_ dimer are within electron transfer distance (Fig. 2c, 3b) [27, 36]. As previously proposed, this could provide an additional source of electrons from H_2_ oxidation by the neighboring dimer subunit to reduce bound O_2_ [36]. However, as Huc operates physiologically under atmospheric conditions, where O_2_ is four million times more abundant than H_2_ [22, 23], it is unlikely that H_2_ oxidation alone would keep the Huc dimers sufficiently reduced to sustainably displace bound O_2_, meaning the enzyme must resist O_2_ inhibition by additional mechanisms.

Concordantly, analysis of the hydrophobic gas channels of Huc that provide H_2_ access to the active site indicates that they are markedly narrower than those of structurally characterized O_2_-sensitive and O_2_-tolerant [NiFe]-hydrogenases (Table S3, Fig 3e, Fig. S8a) [56]. Instead, they are of similar width to the similarly O_2_-insensitive but low-affinity group 1h [NiFe]-hydrogenase from *C. necator* (Table S3) [27, 56]. To determine if these channels play a role in sterically excluding O_2_ from the Huc active site, we performed all-atom molecular dynamics simulations on the HucSL dimer in the presence of excess H_2_ and O_2_ dissolved in water. Strikingly, during these simulations H_2_ enters the Huc active site, while O_2_ is sterically excluded by a series of bottlenecks between the active site and enzyme surface, not coming closer than 5 Å to the catalytic cluster (Fig 3f,g, Fig. S8). The simulations suggest that the most important point selection against O_2_ is a bottleneck after the convergence of the three gas tunnels immediately preceding the active site entrance (Supplemental note 2). This suggests that, as previously proposed for [NiFe] hydrogenases [57], the Huc hydrophobic gas channels play a role in protecting the [NiFe]-cluster from inactivation by O_2_. In turn, the increased selectivity conferred by these gas channels likely decreases the rate of H_2_ diffusion and oxidation compared to O_2_-sensitive and O_2_-tolerant [NiFe]-hydrogenases.

While the narrow hydrophobic gas channels greatly kinetically favour H_2_ diffusion, spectroscopic data indicate that Huc reacts slowly with O_2_. Fourier transform infrared (FTIR) spectroscopy analysis shows that Huc isolated in ambient air exhibits a spectrum consistent with the enzyme adopting the Ni-A or Ni-B state (Fig. 3C, Supplemental note 1). In these O_2_-inhibited states, [NiFe]-hydrogenases bind a bridging hydroxide ligand (μ-OH^−^, Ni-B) or a peroxo ligand (μ-OOH^−^, Ni-A) between the Ni and Fe ions of the catalytic cluster [46, 58, 59]. In the Huc CryoEM structure, additional density is present between the Ni and Fe ions consistent with a single oxygen atom (Fig. 3a), which allows us to assign the IR signature to Ni-B. The assignment of the O_2_ induced state as Ni-B is further supported by the observation that the enzyme readily enters into catalytically active states upon the addition of H_2_ (Fig. S9a,b,c,d, Table S4, Supplemental Note 1). The hydroxide ligand is significantly closer to the Ni ion (1.7-2.0 Å) than the Fe ion (2.2-2.3 Å), indicating a stronger association with the Ni ion (Fig. 3d). There are two plausible mechanisms through which this state can occur: In the direct pathway, assuming O_2_ occasionally reaches the active site, O_2_ can be reduced to two H_2_O with three electrons from the [3Fe-4S] clusters and one from the Ni ion to yield a hydroxide-bound Ni^3+^ species. In the indirect pathway, in line with previous studies showing O_2_-independent formation of Ni-B states [60, 61], oxidation of [3Fe-4S] clusters may lead to oxidation of the [NiFe] centre and a Ni^3+^ ion would be prone to coordinate water with concomitant deprotonation to yield a hydroxide ligand. In either case, Ni-B formation may be protective, given the Ni^3+^ charge in the Ni-B state would decrease the reactivity of the [NiFe] centre to any O_2_ that does reach the active site; by contrast, the kinetically favoured H_2_ ligand would readily displace the hydroxide ligand through a standard ligand substitution reaction. These observations are supported by the kinetics observed in FTIR when reduced Huc is exposed to an atmosphere containing 20% O_2_, with the Ni-S state first populated at the expense of hydrogen bound Ni-R and Ni-C states, before the hydroxide-bound Ni-B state slowly forms (Fig. S9e,f, Supplemental Note 1). In cells, the Ni-B state may occur at lower rates, due to the enzyme environment and menaquinone-dependent redox control.

The high-affinity kinetics of Huc differ radically from more classical O_2_-sensitive hydrogenases, which tend to be faster low-affinity enzymes [5, 62]. A comparison of the environment surrounding the [NiFe] active site of Huc and O_2_-sensitive hydrogenases reveals no differences that could readily explain the divergent properties of these enzymes (Fig. S10a,b,c). This indicates that the overall activity of the [NiFe] active site instead results from changes in the properties of the iron-sulfur clusters of the small subunit, or other regions of the enzyme. Given their distinct structure, the modified gas channels identified in Huc may play a role in selectively capturing H_2_ molecules and delivering them to the active site, decreasing the rate but increasing the affinity of the enzyme [57, 63]. In addition, the increased redox potential of Huc (Eon ≈ −140 mV) means that atmospheric H_2_ oxidation, with a redox potential (E_atm_) of ≈ −0.134 mV at pH 7, is thermodynamically favourable in comparison to low-affinity variants operating at minimal overpotential (Eon ≈ −0.36 V) [64]. The three [3Fe-4S] clusters are likely the main determinants of the high potential of the enzyme [48, 65]. Moreover, the redox properties of the proximal and medial clusters may be further modified by the presence of an unusual D-isomer of histidine at position 166 of HucL (Fig. S10d) [66]. An L-isomeric histidine is conserved at this position in other characterized hydrogenases, with its imidazole head group interacting with the proximal cluster (Fig. S10e). In Huc, His-166 also interacts with the proximal cluster (Fig. 3h), but the D-isomeric configuration shifts the position of the backbone carbonyl of His-166 so that it interacts with a highly coordinated ligand in proximity to the medial cluster (Fig. 3h, Fig. S10f). We were unable to identify this ligand, though its elongated density suggests it is not water and it may play a role in tuning the redox potential of the medial [3Fe-4S] cluster. The key determinants of the high affinity of Huc require further experimental investigation, for which the structure of Huc will be of critical importance for hypothesis generation and experimental design.

### Huc extracts, transports, and directly reduces menaquinone from the cell membrane in an internal hydrophobic chamber

Analysis of the Huc CryoEM electron density maps reveals additional electron density near the distal [3Fe-4S] cluster of each Huc protomer (Fig 4ai). Using maps from the high-resolution reconstruction, this electron density could be unambiguously modelled as the redox-active head group of menaquinone (Fig. 4aii). In the lower resolution oligomer maps, weaker electron density is also present for part of the aliphatic menaquinone tail (Fig. 4aiii). The head group of menaquinone is stabilized by hydrogen bonding between its oxygen groups and the terminal hydroxyl of Tyr-301 and the backbone carbonyl of Lys-212 of HucS, as well as π-π interactions with Tyr-229 of HucS (Fig 4bi). Interestingly, in our 1.67 Å reconstruction, Tyr-229 adopts a second conformation that clashes with density corresponding to menaquinone (Fig. 4bii, Mov. S3). In this conformation, Tyr-229 shields the proximal [3Fe-4S] cluster from the solvent and may assist in preventing the Huc electron transport chain from reducing O_2_ in the absence of the menaquinone substrate, which would lead to the formation of reactive oxygen species. Confirming the assignment of this density as menaquinone, mass spectrometry detected high levels of beta-dihydromenaquinone-9 associated with purified Huc (Fig. 4c). This indicates this location is the electron acceptor substrate binding site of Huc and that the enzyme directly binds and reduces menaquinone. This observation is consistent with our enzyme activity data showing that menadione is the preferred electron acceptor for Huc (Fig. 1c). These findings contrast with observations made in group 1 [NiFe]-hydrogenases, where small subunits are electron relays without catalytic roles [24–28].

**Figure 4.**
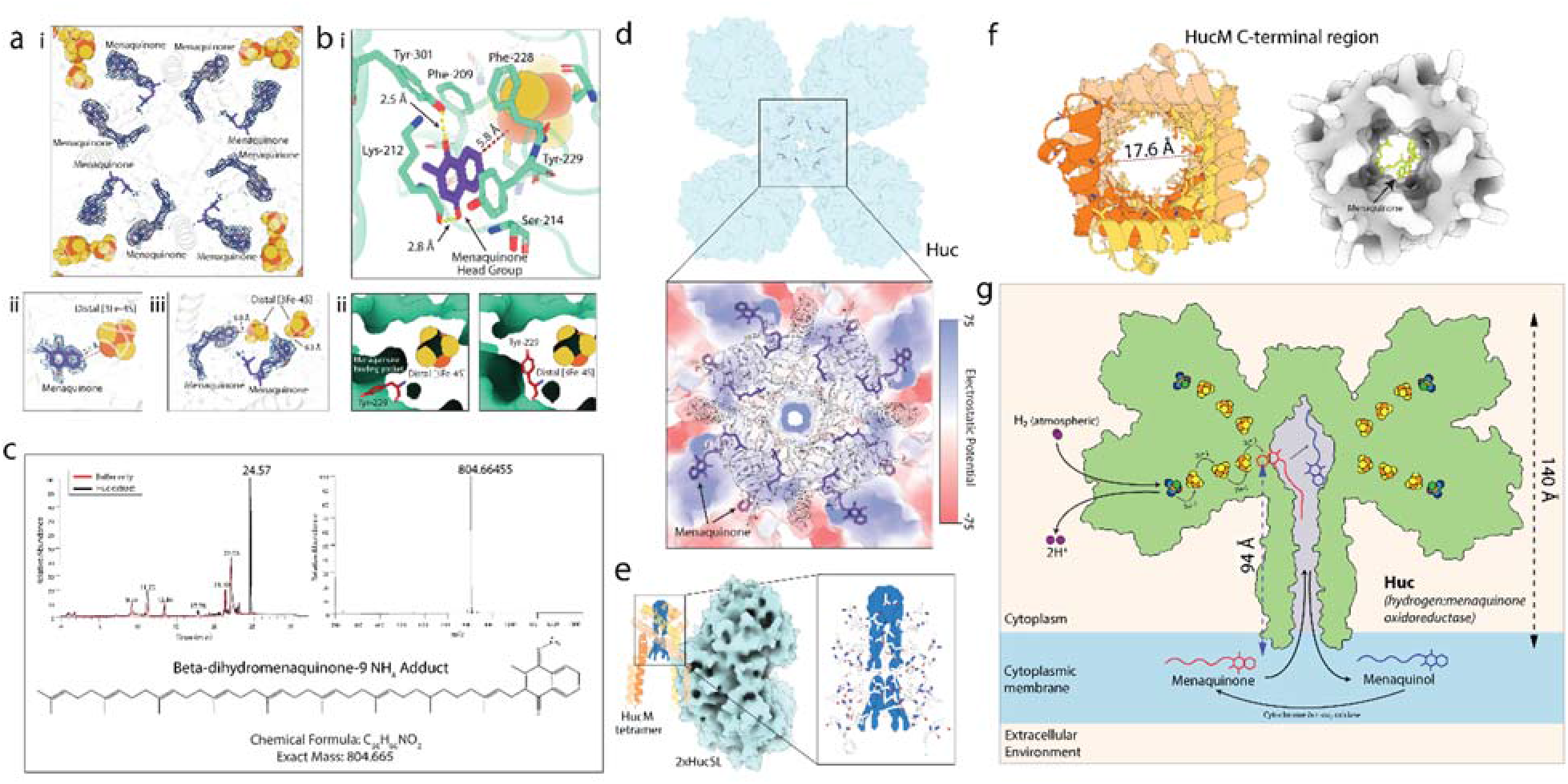
Huc extracts menaquinone from the membrane and directly reduces it. (a-i) A view of the chamber at the centre of the Huc oligomer showing electron density (blue mesh) resulting from bound menaquinone from the lower resolution Huc oligomer CryoEM reconstruction. (a-ii) Electron density for bound menaquinone in the high-resolution Huc dimer CryoEM maps. Electron density for the menaquinone tail is absent in these maps due to symmetry averaging. (a-iii) A zoomed view of menaquinone molecules associated with a lobe of the Huc oligomer, showing the distance of each molecule to the distal [3Fe-4S] cluster. (b-i) The coordination of the menaquinone head group by the HucS subunit, within electron transfer distance of the distal [3Fe-4S] cluster. (b-ii) The two conformations observed for Tyr-229 of HucS in the high-resolution Huc CryoEM electron density maps. Tyr-229 adopts an open conformation in the presence of menaquinone (left panel) and a closed conformation (right panel) that is mutually exclusive to menaquinone binding. (c) (c) HPLC-MS analysis of the Folch extract from purified Huc. The base peak chromatogram (left panel) shows a substantial peak at 24.6 min corresponding to an ion at m/z = 804.66455 (right panel), which is consistent with the ammonium adduct of beta-dihydromenaquinone-9, m/z = 804.66531 (bottom panel) (d) A hybrid stick and electron static surface representation of the central cavity Huc, showing the surface of the chamber is lined with hydrophobic residues. (e) A view of the HucM tetramer showing the presence of electron density for lipids that occlude the top of the Huc internal hydrophobic chamber. (f) A top-down view of the HucM C-terminal tube, showing the interior of the tube is lined with hydrophobic residues (left) and is capable of accommodating at menaquinone molecule (right). (g) A model for reduction of menaquinone by Huc, using electrons derived from the oxidation of atmospheric H_2_.

However, despite this strong evidence that Huc directly donates electrons from H_2_ oxidation to respiratory quinone, Huc is not an integral membrane protein and thus the highly hydrophobic, membrane-bound menaquinone must be transported to its substrate-binding site 94 Å away from the membrane surface. Analysis of the structure of the Huc oligomer indicates the mechanism by which this occurs. The interior surface of the internal cavity of Huc is lined by hydrophobic sidechains, contributed by both the HucS and HucM subunits (Fig. 4d, Mov. S4). Thus, the interior surface of the chamber is suited to the binding of menaquinone and other hydrophobic molecules. Consistently, the top of the Huc chamber contains cylinders of electron density, coordinated by hydrophobic sidechains, which likely correspond to lipids, occluding this entrance of the chamber (Fig. 4e). The conduit for menaquinone entry to and exit from the hydrophobic chamber is provided by the membrane-associated Huc stalk. The hydrophobic interior of the HucM tube that constitutes this stalk has dimensions capable of accommodating menaquinone (Fig. 4f, Fig. S7d), which can enter through the interaction of HucM with the cell membrane (Mov. S4). Menaquinone molecules likely diffuse through this tube into the Huc hydrophobic chamber, where they bind the substrate acceptor site and are reduced to menaquinol by electrons derived from atmospheric H_2_ oxidation. Reduced menaquinol then diffuses back into the membrane via the HucM C-terminal tube, where it is oxidized selectively by the cytochrome *bcc-aa3* oxidase terminal oxidase to generate proton motive force (Fig 4g) [32]. Among hydrogenases, this mechanism of quinone reduction is unique to Huc, with the other structurally characterised respiratory hydrogenases reducing quinone in the membrane via direct electron transfer to an integral membrane cytochrome *b* subunit [67, 68]. There are, however, some similarities with the reduction mechanism of respiratory chain complex I, which removes quinone ~30 Å from the membrane to position it for reduction by a [4Fe-4S] cluster within its soluble arm; however, in complex I [69, 70], this extraction is achieved via an enclosed channel, rather than the broad hydrophobic tube and chamber observed in Huc [69].

## Discussion

To use the trace quantities of H_2_ present in air, the [NiFe]-hydrogenases of trace gas scavenging bacteria require distinct properties compared to their counterparts that function under H_2_-rich hypoxic or anoxic conditions [5, 46]. Through biochemical and electrochemical characterisation of Huc from the aerobic bacterium *M. smegmatis,* we show that the required properties of O_2_ insensitivity and high affinity for H_2_ are inherent to this hydrogenase, rather than resulting from coupling to other processes inside the bacterial cell. Moreover, through determination of the CryoEM structure of Huc, MD simulations, and FTIR spectroscopy, we provide strong evidence that at least partial exclusion of O_2_ from the active site contributes to O_2_ insensitivity of the enzyme. Our data shows that Huc has a large electrochemical overpotential that makes it uniquely tuned for oxidation of trace quantities of H_2_ and for the direct donation of the resulting electrons to the respiratory cofactor menaquinone. Strikingly, we demonstrate that Huc accesses menaquinone via a unique and highly unusual mechanism. Through the scaffolding protein HucM, the Huc complex can extract menaquinone from the membrane and transport it 94 Å to the enzyme’s electron acceptor site. This finding greatly expands our knowledge of the possibilities for performing respiratory quinone reduction. More broadly, these findings open pathways for biocatalyst development given all hydrogenases applied in whole-cell and purified enzyme systems are low-affinity enzymes that become inhibited O_2_. Huc, as an oxygen-insensitive high-affinity enzyme and the first structurally characterised group 2 [NiFe]-hydrogenase, provides a basis for the development of biocatalysts that operate under ambient conditions.

## Acknowledgments

We thank the Bio21 Institute for the use of the Thermo Glacios housed in their facilities during CryoEM sample preparation and screening. We thank Prof. Jamie Rossjohn, Prof. Fraser Armstrong, and Dr. Gavin Knott for their review and constructive feedback on the manuscript.

## Funding

Australian Research Council DECRA Fellowship (DE170100310) (CG)

Australian Research Council (DP200103074) (RG, CG)

National Health & Medical Research Council Emerging Leader Fellowships (APP1178715) (CG)

National Health & Medical Research Council Emerging Leader Fellowships (APP1197376) (RG)

Deutsche Forschungsgemeinschaft through Priority Program 1927 (1554/5-1) (STS)

The Swedish Energy Agency (grant no 48574-1) (GB)

The European Research Council (714102) (GB)

European Union’s Horizon 2020 research and innovation program (Marie Skłodowska Curie Grant No. 897555 (MS)

## Author contributions

Conceptualization: CG, RG, AK, PC, GMC

Methodology: RG, AK, CG, HV, JB, MS, PC, CKB, SK, GB

Investigation: RG, AK, PC, HV, JB, MS, PC, STS, KAV, CKB, MB, CG

Visualization: RG, AK, HV, JB, MS, PC, CKB

Funding acquisition: CG, RG, GB, MS

Project administration: CG, RG

Supervision: CG, RG, GB, KS, RBS, SK

Writing – original draft: RG, CG, AK

Writing – review & editing: RG, CG, AK, all other authors

## Data and materials availability

Cryo-EM maps and atomic models generated in this paper have been deposited in the Electron Microscopy Data Bank (accession codes 7UTD, 7UTX, 7UUR, 7UUS) and the Protein Data Bank (accession codes EMD-26767, EMD-26793, EMD-26801, EMD-26802). Other source data are provided with this paper. Prior to submission, the resolution of the map associated with 7UTX was improved from 1.57 Å to 1.52 Å, this higher resolution data was being processed by the PDB at the time of manuscript submission. We are experiencing extended delays (>2 weeks) in receiving final validation reports from the PDB, so in order not to delay submission, the final validation reports from the 1.57 Å reconstruction and a draft validation report for the improved 1.52 Å reconstruction are provided.

## Supplementary Text

Figs. S1 to S10

Tables S1 to S5

Movies S1 to S4

Supplemental Notes 1 and 2

## Supplemental Figures

**Figure S1.**
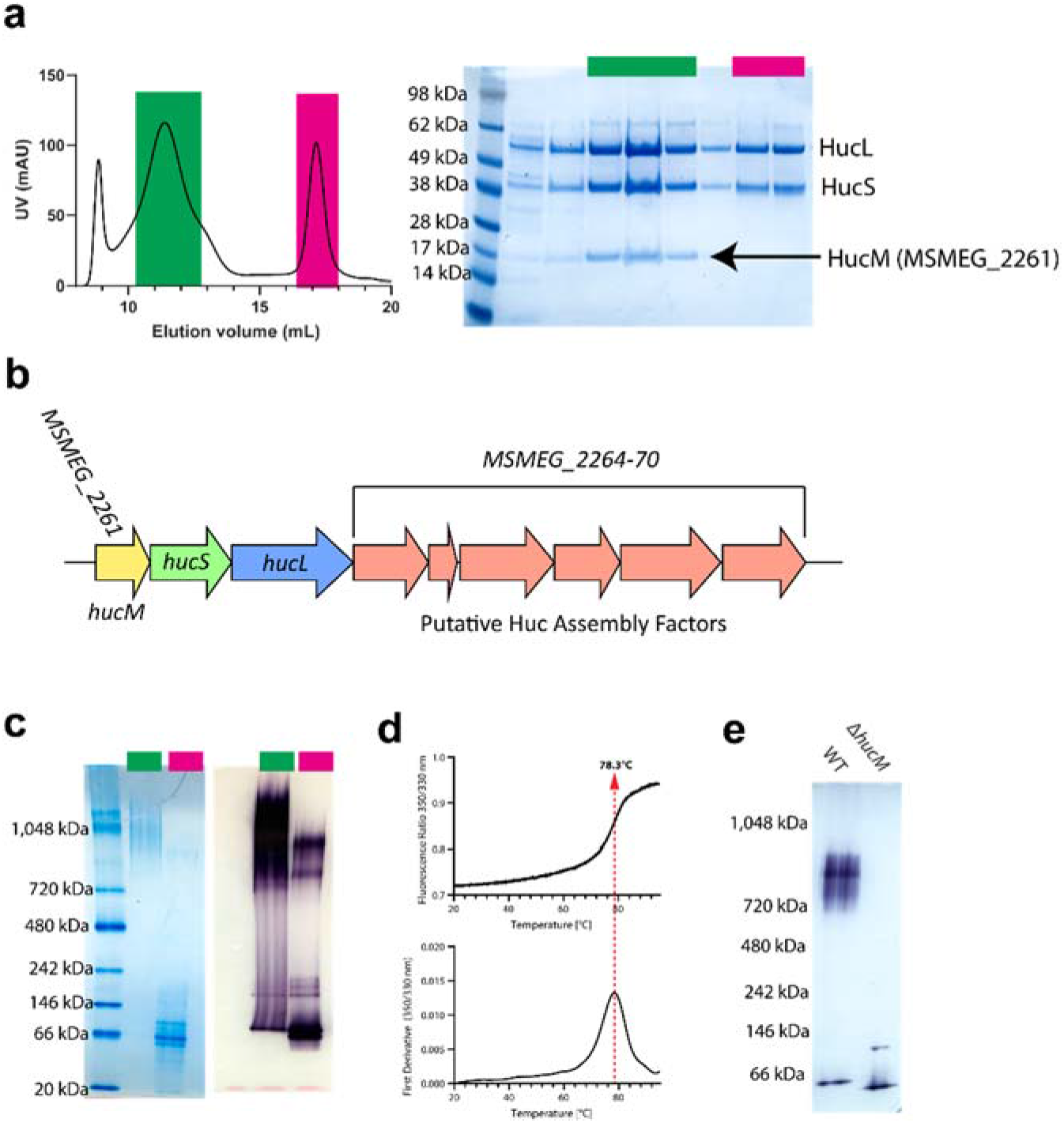
Huc isolation and purification. (a) Left panel: A chromatogram showing streptactin purified Huc separated via size exclusion chromatography on a Superose 6 10/300 column. The green highlighted region contains the Huc oligomer, while the pink region contains a low molecular weight Huc species. Right panel: A coomassie stained SDS-PAGE gel showing fractions from the coloured peak regions of the chromatogram. (b) A schematic of the Huc gene cluster showing the location of *hucM* (MSMEG_2261) compared to *hucS* and *hucL.* (c) A native-PAGE gel of the purified Huc oligomer (green) and low molecular species (pink), stained with coomassie (left panel) and NBT (right panel). (d) Differential scanning calorimetry analysis of the Huc oligomer showing a transition in tryptophan derived fluorescence at 78.3°C, indicating the melting temperature of the oligomer. (e) An NBT stained native-PAGE gel of *M. smegmatis* cell lysate showing that Huc does not form an oligomer in the Δ*hucM* strain.

**Figure S2.**
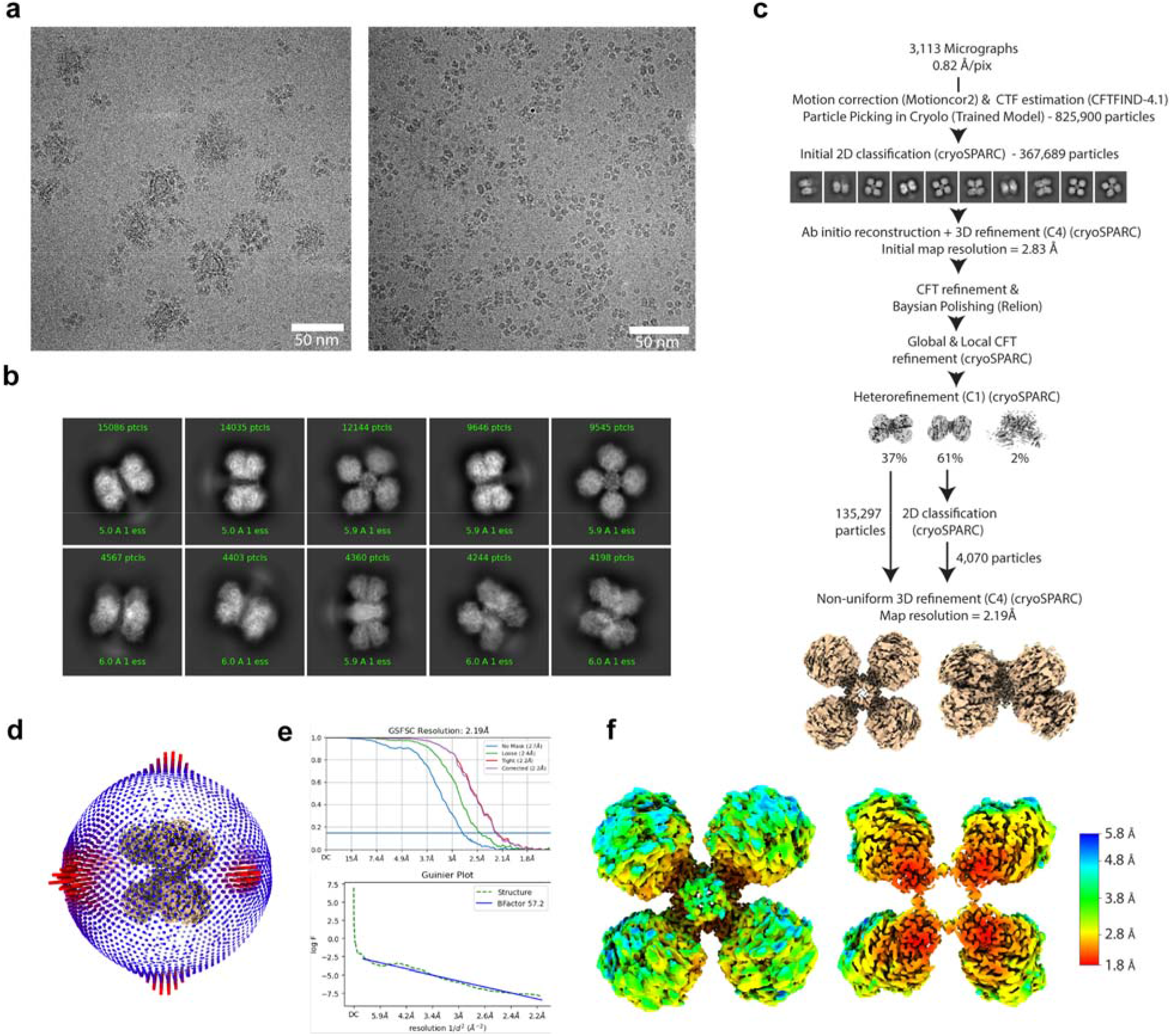
CryoEM visualisation and 3D reconstruction of the Huc oligomer. (a) motion-corrected micrographs of vitrified purified Huc oligomers, showing free Huc (right panel) and Huc associated with membrane vesicles (left panel). (b) Selected 2D class averages of Huc oligomer, showing C4 symmetry and flexible membrane-associated stalk. (c) Data processing workflow for the Huc oligomer reconstruction. (d) The Euler angle distribution of particles used for Huc oligomer reconstruction. (e) Gold-standard Fourier shell correlation (FSC) curves calculated from two independently refined half-maps, indicate an overall resolution of 2.19 Å at FSC = 0.143, and Guinier plot indicates a sharpening B-factor of 57.2. (f) A local resolution map of the Huc oligomer, showing a resolution range of ~1.8 to 4.8 Å from the Huc core to the periphery indicative of significant interdomain motion.

**Figure S3.**
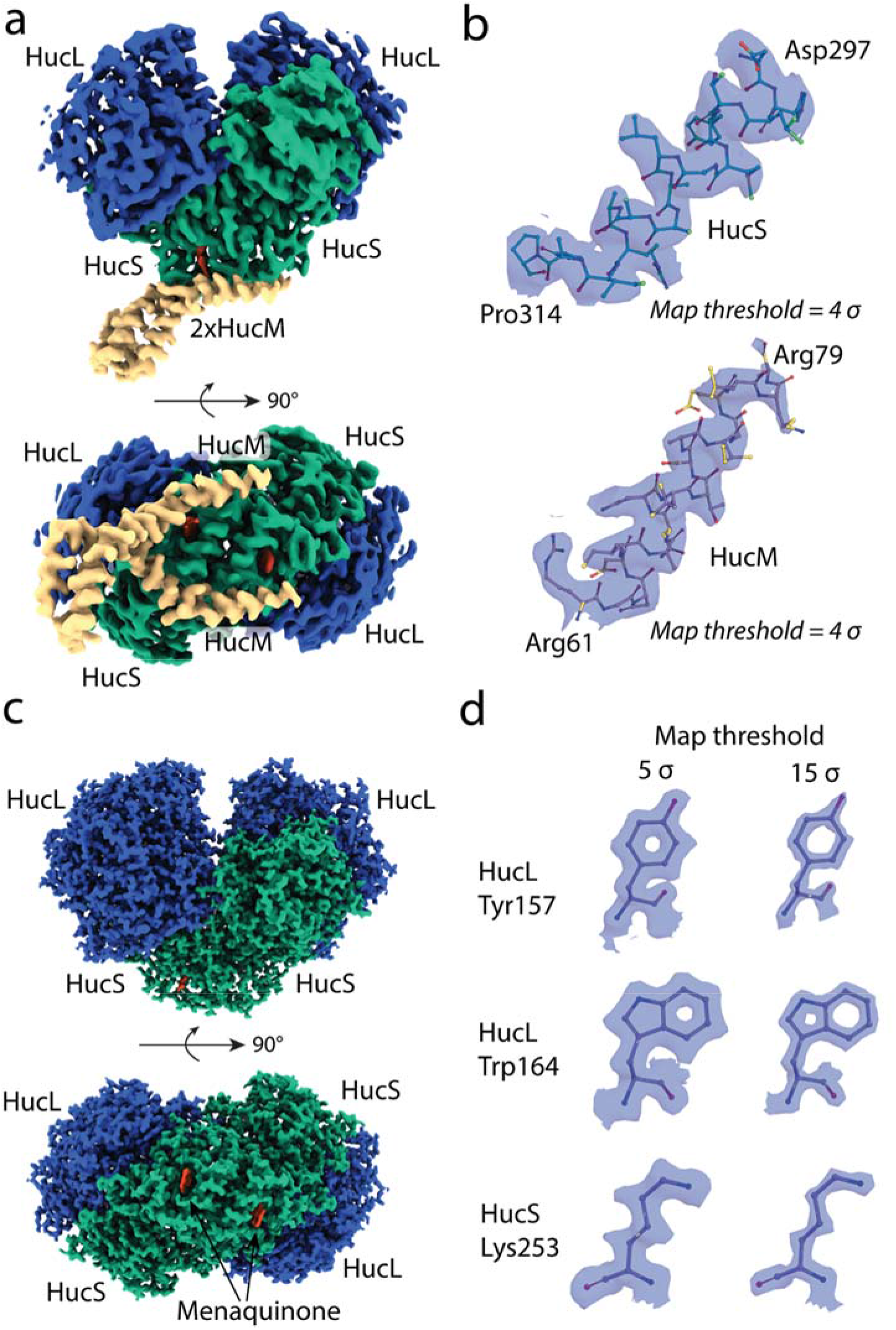
Map quality of Huc oligomer and dimer CryoEM reconstructions. (a) Maps from the 2.19 Å Huc oligomer reconstruction for one Huc lobe (HucS_2_L_2_) and 2 HucM molecules. (b) Helices from HucS and HucM with associated electron density from oligomer reconstruction. (b) Maps from the 1.52 Å Huc dimer reconstruction. (d) Examples of electron density corresponding to amino acids from the Huc dimer reconstruction contoured at 5 or 15 σ.

**Figure S4.**
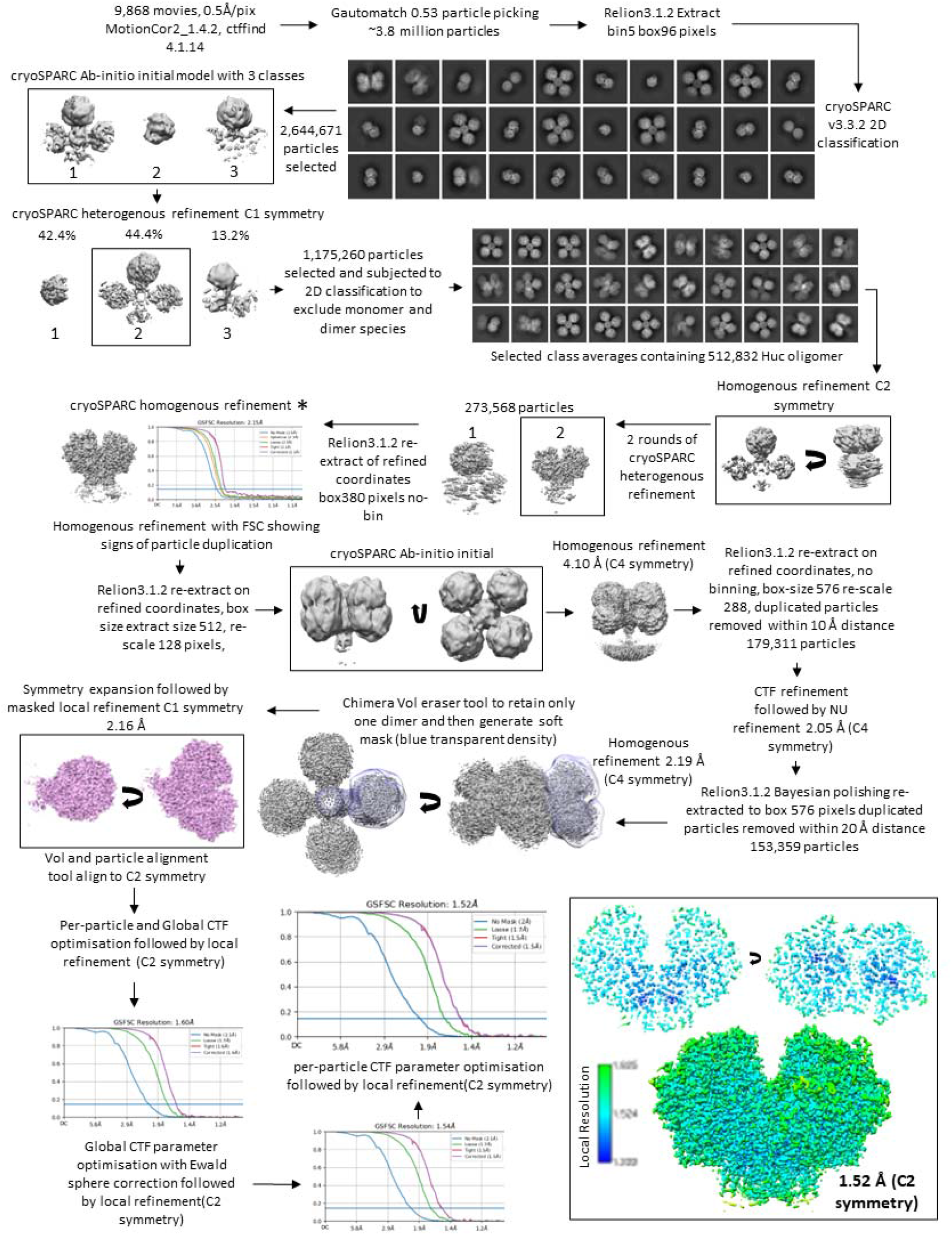
Data processing workflow for the 1.52 Å Huc dimer reconstruction.

**Figure S5.**
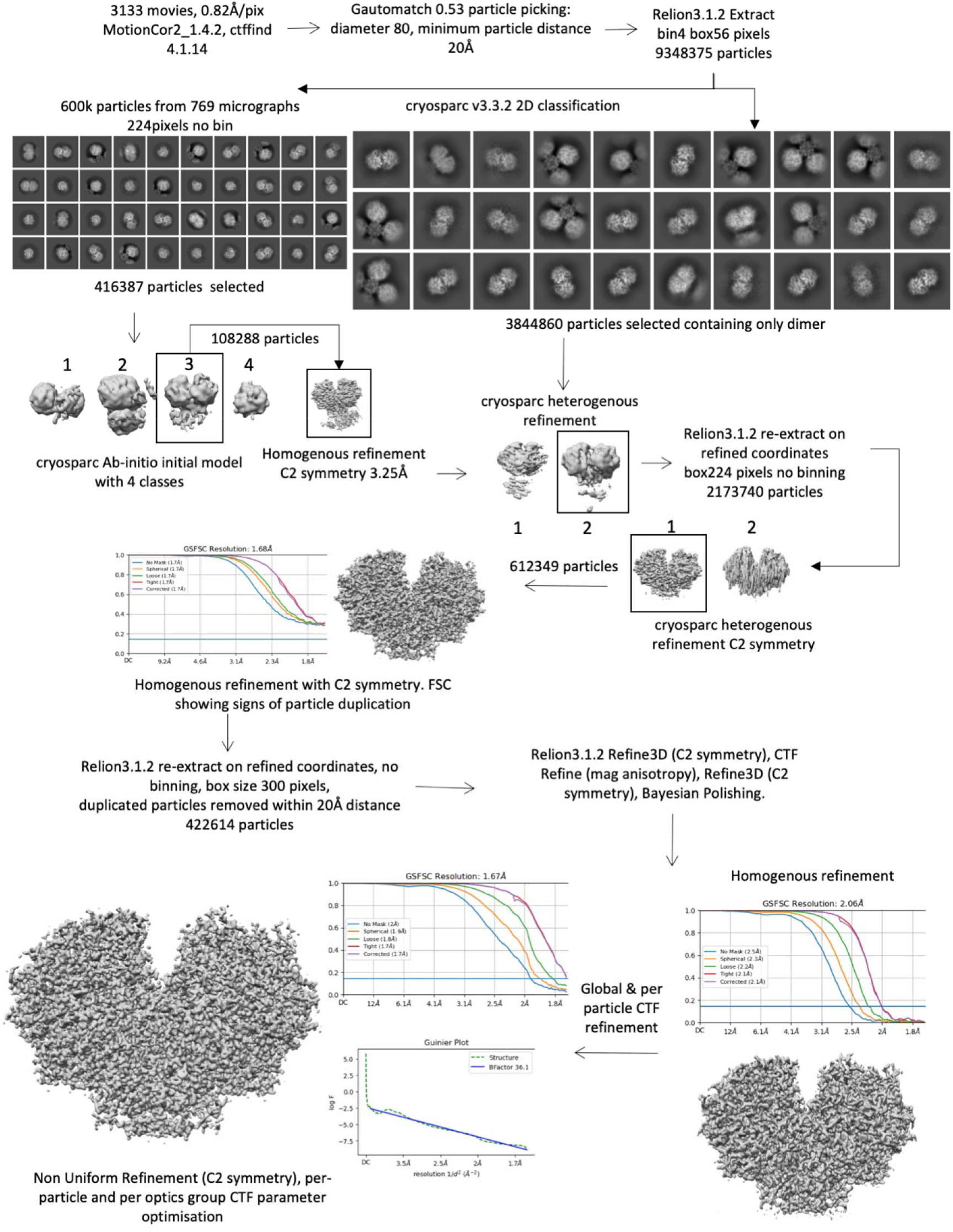
Data processing workflow for the 1.67 Å Huc dimer reconstruction.

**Figure S6.**
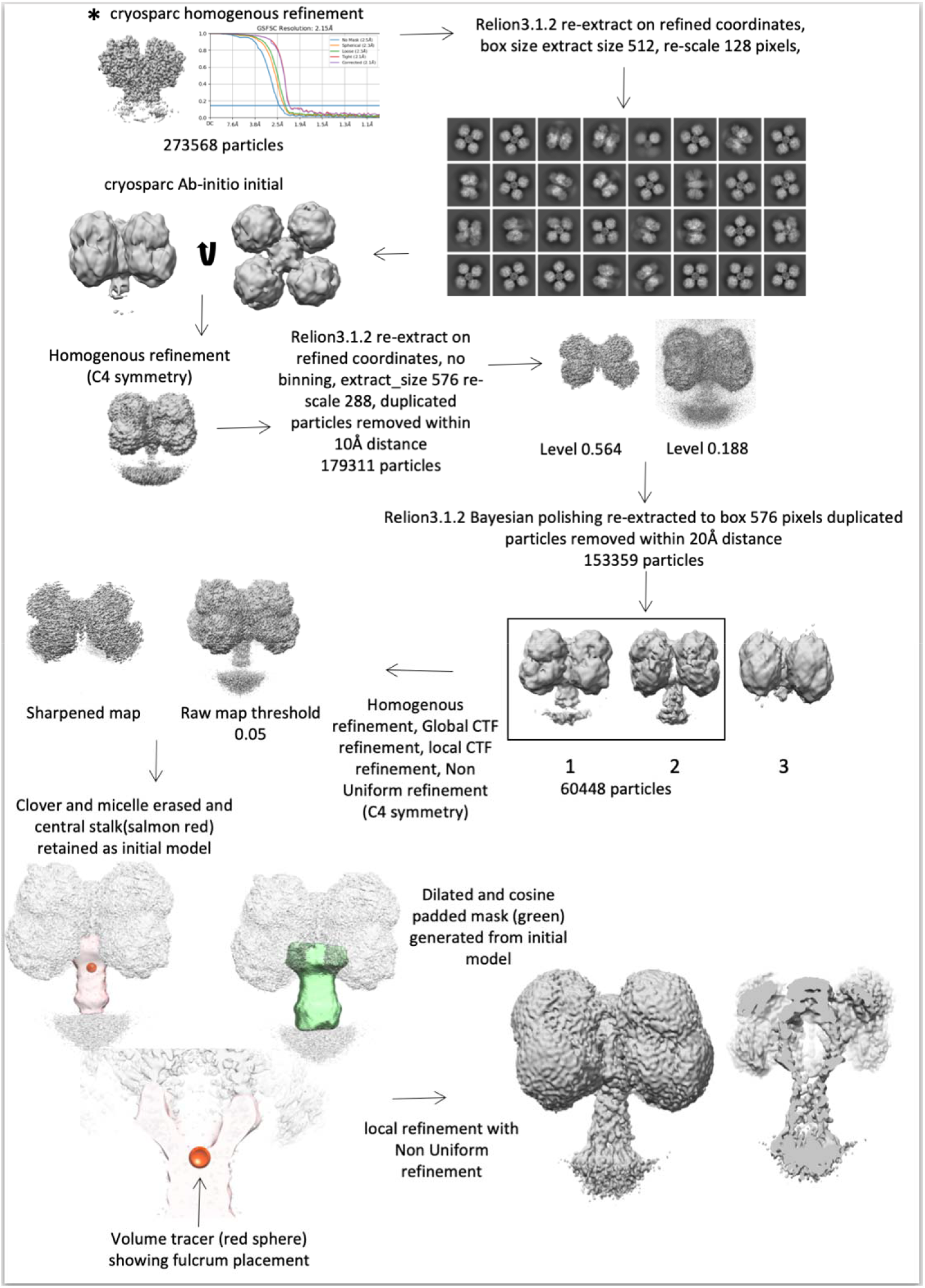
Data processing workflow for Huc stalk reconstruction. * corresponds to particles from homogenous refinement indicated in Fig. S4.

**Figure S7.**
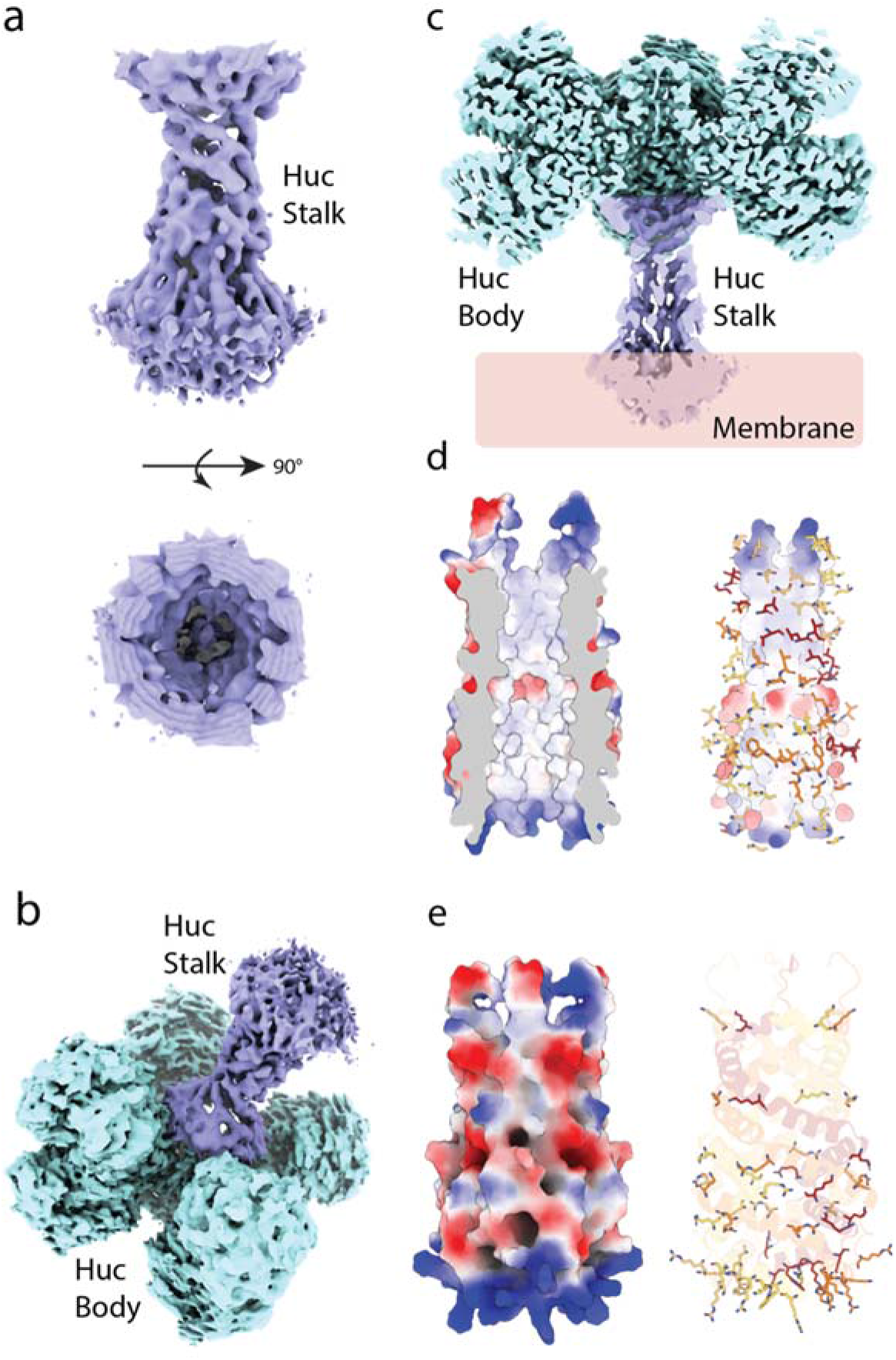
CryoEM electron density map and AlphaFold model of the Huc stalk region. (a) The stalk region from electron density maps from Fig. S6. (b) A composite of the Huc stalk and body map regions from the 2.19 Å dataset. (c) A cut-away view of the composite map from panel b, showing the enclosed chamber and the centre of the Huc complex. (d) A surface view of the AlphaFold model of the HucM c-terminal stalk region, shown as a cutaway electrostatic surface (left) and the internal surface lined with hydrophobic residues (right). (e) The HucM c-terminal region as in panel d showing the positively charged region at the base of the stalk (left) and the high frequency of arginine and lysine residues in this region (right).

**Figure S8.**
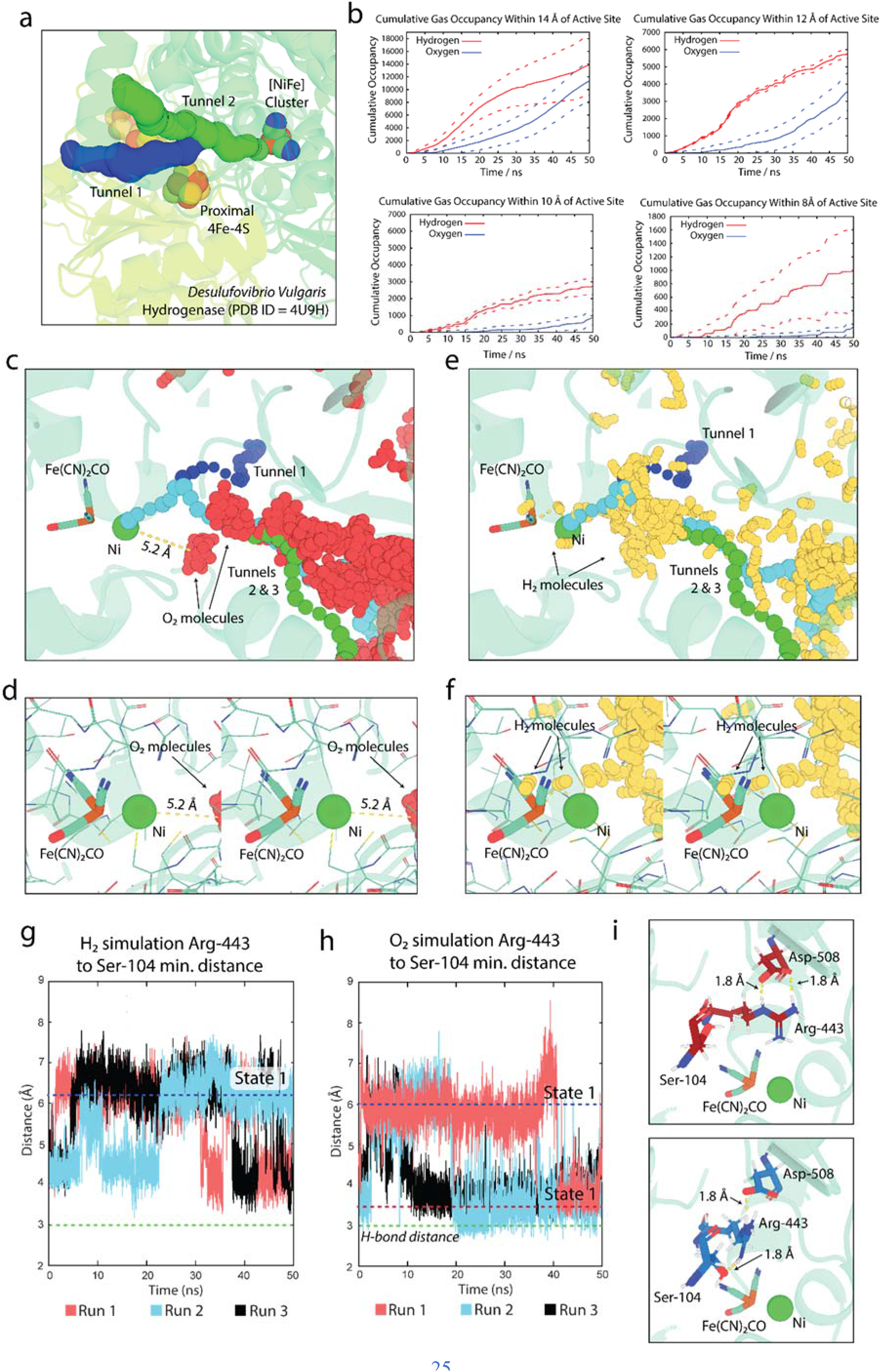
Molecular dynamics simulations of HucSL in the presence of H_2_ or O_2_. (a) A zoomed view of the large and small subunits of an [NiFe]-hydrogenase from *Desulfovibrio vulgaris* (PDB ID = 4U9H) [28], showing the location and width of the hydrophobic gas channels that provide substrate access to the [NiFe] active site. See Table S3 for tunnel width statistics. (b) Cumulative occupancy plots showing the proximity of H_2_ and O_2_ molecules to the [NiFe] cluster of Huc throughout the molecular dynamics simulations. (c) An expanded view of the positions of a representative subset of O_2_ molecules in closest proximity to the Huc [NiFe] cluster in molecular dynamics simulations, the path of the hydrophobic gas channels is shown as blue, green, and cyan spheres. (d) A zoomed stereo view of O_2_ as described in panel c. (e) and (f) The positions of a representative subset of H_2_ molecules in closest proximity to the Huc [NiFe] cluster are displayed as described in panels c and d. (g) A distance-over-time plot showing the relative proximity of serine 104 to arginine 443, in the Huc MD simulations in the presence of H_2_. (h) A distance-over-time plot showing the relative proximity of serine 104 to arginine 443, in the Huc simulations in the presence of O_2_. (i) The relative position of arginine 443 in states 1 (top panel) and 2 (bottom panel) populated during the Huc MD simulations.

**Figure S9.**
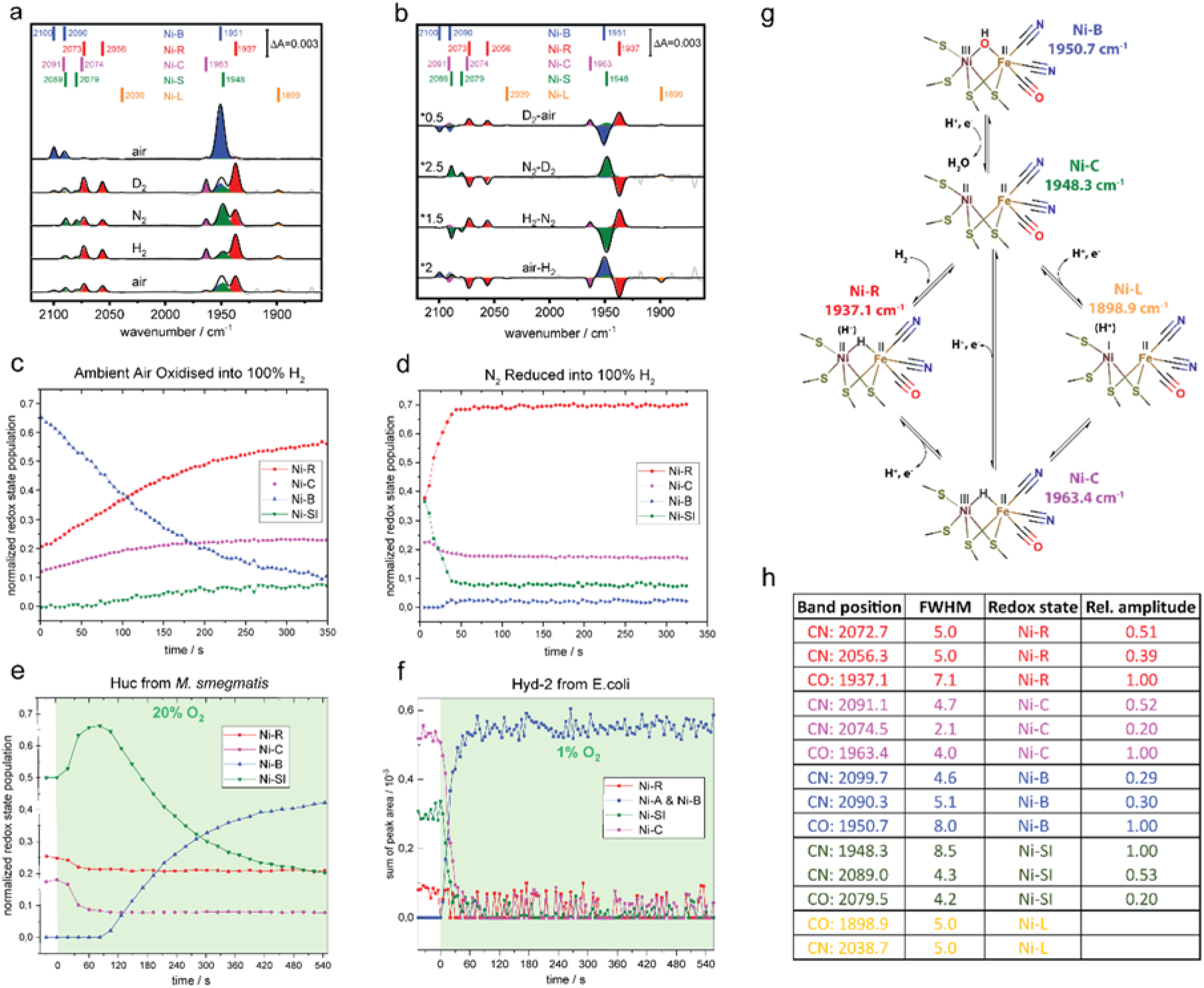
FTIR analysis of Huc in the presence of ambient air, N_2_ or H_2_. (a) Absolute FTIR spectra of Huc equilibrated in air, followed by sequential equilibration in a 100% atmosphere of the labelled gas, followed by air. [NiFe] cluster states are assigned based on literature-derived values (see Table S4). (b) Difference spectra derived from spectra shown in panel a. (c) A time course plot showing the relative fractions of the FTIR assigned states immediately following the transfer of Huc from ambient air to a 100% H_2_ atmosphere, the catalytic Ni-R state becomes populated at the expense of hydroxide bound Ni-B over a time scale of minutes. (d) A time-course plot showing the relative fractions of the FTIR assigned states immediately following the transfer of Huc in a mixed Ni-S, Ni-R state from 100% N_2_ to a 100% H_2_ atmosphere. The catalytic Ni-R state becomes populated at the expense of the non-hydrogen bound reduced Ni-S state over a timescale of seconds. (e) A time-course plot of [NiFe] cluster states following the transfer of Huc predominantly in the Ni-S state from 100% N_2_ into 80%:20% N_2_ to O_2_. Initially, the Ni-S state is populated at the expense of the hydrogen bound Ni-C and Ni-R states, followed by population of the Ni-B state at the expense of Ni-S over a timescale of minutes. (f) A time-course plot of [NiFe] cluster states of Hyd-2 from *E. coli* following the transfer from 100% N_2_ into 99%:1% N_2_ to O_2_. The Ni-B state is rapidly populated at the expense of all other states over a timescale of seconds. Data from Senger et al. 2018 [71]. (g) A scheme of the proposed catalytic cycle for [NiFe]-hydrogenases, showing the states occupied by Huc during FTIR analysis. (h) A table of the wavenumber of the peaks assigned to the CO band of states identified in the Huc FTIR spectra.

**Figure S10.**
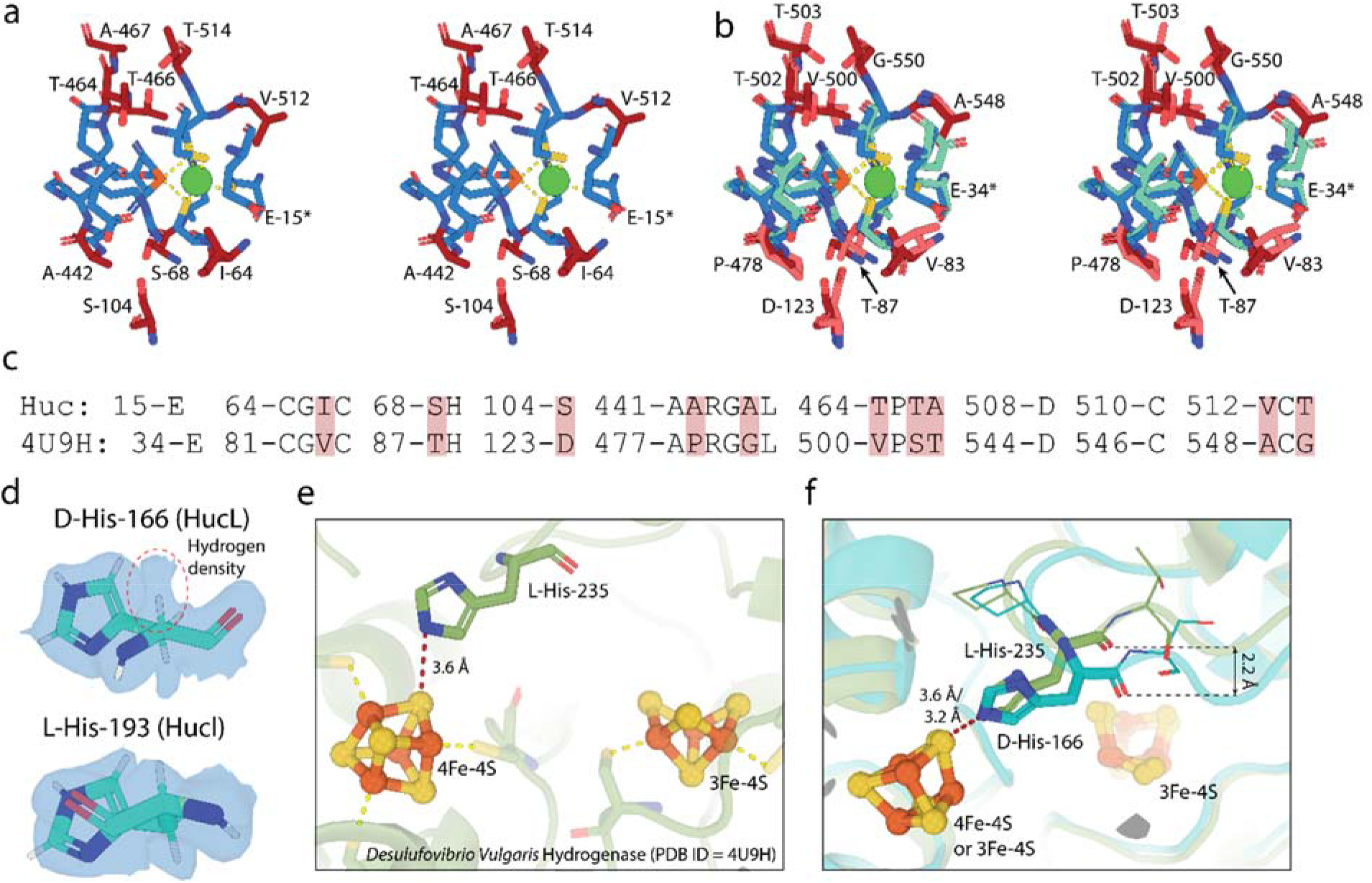
Comparison of Huc with an O_2_ sensitive [NiFe]-hydrogenase. (a) A stereoview of the Huc [NiFe] cluster and surrounding amino acids. Amino acids conserved in O_2_ sensitive hydrogenases and shown in blue, while divergent residues are shown in red. Amino acid type and position are labelled for divergent residues. (b) A stereoview overlay of the Huc catalytic site residues from panel a with those of the O_2_ sensitive hydrogenase from *D. vulgaris* (PDB ID = 4U9H). Huc residues are coloured as in panel a, while *D. vulgaris* hydrogenase residues are coloured cyan (conserved) and pink (divergent). (c) A sequence alignment of Huc and *D. vulgaris* hydrogenase catalytic site residues, with divergent residues highlighted in red. (d) A stick view of D-histidine 166 from HucL showing amino acid chirality, the corresponding electron density map is shown at 5 σ, demonstrating the density is consistent with the D-isomer (top panel). L-histidine 194 form HucL is shown for reference (bottom panel). (e) The location of L-histidine 235 from the *D. vulgaris* hydrogenase, which corresponds to D-histidine 166 from Huc, showing its interaction with the proximal 4Fe-4S cluster. (f) An overlay of the relative positions of D-histidine 166 from Huc and L-histidine 235 from the *D. vulgaris* hydrogenase showing how the D-isomer shifts the position of the mainchain in Huc.

## Supplemental Tables

**Table S1 Huc H_2_ oxidation kinetic parameters**

**Table S2 Huc CryoEM data collection, processing, and refinement statistics**

**Table S3 Caver generated hydrogenase gas channel statistics**

**Table S4: Comparison of Huc FTIR spectral states with previously characterised hydrogenases (adapted from [72]) (units = cm^−1^)**

**Table S5: Bacterial strains and plasmids used in this study**

## Supplemental Movies

**Movie S1 3D variability analysis of Huc reveals significant inter-lobe movement.**

**Movie S2 Reconstruction of the structure of the full-length Huc complex.**

**Movie S3 The Huc menaquinone binding site is gated by Tyr-229 of HucS**

**Movie S4 Huc possesses an internal hydrophobic chamber for reducing menaquinone**

## Supplemental Notes

**Supplemental Note 1. Analysis of Huc redox states by aerosol-controlled ATR-FTIR spectroscopy**

To gain an understanding of the redox states adopted by the Huc [NiFe] cluster we recorded its FTIR spectra sequentially in ambient air, H_2_, and N_2_. Huc prepared under ambient conditions (78.09% N_2_, 20.95% O_2_, 0.000053% H_2_) was analysed under 100% N_2_. Under these conditions, the FTIR spectra were assigned to the oxidized Ni-B state, which possesses an OH^−^ ligand bridging the catalytic Ni and Fe ions, since the CryoEM structure obtained under similar conditions shows only electron density for one oxygen atom, which renders the formation of the Ni-A state with a peroxo ligand (OOH^−^) unlikely (Fig. S9a,b). This indicates that the Huc [NiFe] cluster is fully oxidized by the O_2_ present in ambient air. When the atmosphere in the cell was exchanged for ^2^H_2_, the Ni-B signal diminished and spectra consistent with a high proportion of the catalytic Ni-R and Ni-C state, and a lesser amount of Ni-S state were observed (Fig. S9a,b,g,h). This indicates that the oxidized Ni-B state of Huc can rapidly enter the H_2_ oxidation cycle when H_2_ becomes available. The subsequent exchange of the cell atmosphere for N_2_ led predominantly to population of the Ni-S state, a reduced, non-hydrogen bound state possibly through auto-oxidation by Huc-associated menaquinone (Fig. S9a,b). A further exchange of the atmosphere with H_2_ led to a similar spectrum to that observed for ^2^H_2_ (Fig. S9a,b). Finally, when the atmosphere was exchanged for air, the proportion of the Ni-B state increased at the expense of the reduced states, confirming the role of O_2_ in oxidizing the enzyme to this state (Fig. S9a,b). Taken together, these data indicate that like O_2_-tolerant membrane-bound hydrogenases, the Huc [NiFe] cluster oxidises to the hydroxyl bound Ni-B state in the presence of O_2_ [46, 73, 74]. To better understand the kinetics of the redox state transitions of the Huc [NiFe] cluster we collected time-resolved FTIR spectra after transferring Huc oxidised in ambient air into a 100% H_2_ atmosphere. Over a timescale of minutes, the Ni-R and Ni-C states were populated at the expense of Ni-B with a near-equilibrium reached by 350 seconds (Fig. S9c). In contrast, when reduced Huc populating predominantly Ni-R and Ni-S states was transferred from 100% N_2_ into 100% H_2_ it rapidly populated the Ni-R state at the expense of Ni-S, equilibrium was achieved within 50 seconds (Fig. S9d). These data indicate that Huc in the hydroxide bound Ni-B state reacts more slowly with H_2_ than in the reduced non-hydrogen Ni-S bound state. This is consistent with the [NiFe] bound hydroxyl in the Ni-B state needing to be displaced by H_2_ for the catalytic cycle to proceed. To assess the rate response of the Huc [NiFe] cluster when exposed to O_2_, we collected time-resolved FTIR spectra after transferring Huc in a mixed Ni-S, Ni-R, and Ni-C state from 100% N_2_ into 20%:80% N_2_ to O_2_. Initially, the Ni-S state was further populated at the expense of Ni-R and Ni-C, before the Ni-S state was replaced by Ni-B over a timescale of minutes (Fig. S9e). These data suggest that the hydrogen-bound Ni-R and Ni-C states must first convert to the empty reduced Ni-S state before the hydroxide-bound oxidised Ni-B state is formed (Fig. S9g,h). This is in contrast to the oxidation profile observed for the oxygen-sensitive hydrogenase Hyd-2 from *E. coli* which was previously shown to rapidly adopt Ni-A and Ni-B states at the expense of all other states when exposed to only 1% O_2_ (Fig. S9f) [71]. These data indicated that the Huc [NiFe] cluster is oxidised much more slowly than oxygen-sensitive hydrogenases, likely due to the restricted access of O_2_ to its active site and potentially because of the altered properties of its redox chain.

**Supplemental Note 2. MD simulations of H_2_ and O_2_ diffusion towards the Huc active site**

To test the hypothesis that narrowing of the hydrophobic tunnels that provide access to the Huc active site contributes to the O_2_ insensitivity of the enzyme, we performed molecular dynamics simulations on the HucSL dimer in the presence of an excess of either H_2_ or O_2_. Simulations were run for 50 nanoseconds (ns) and repeated three times. Huc was stable and retained its original secondary structure throughout the simulations. H_2_ penetrated the Huc gas channels, with the molecule reaching within 8 Å of the active site within the first 10 ns of the simulations (Fig. S8b). O_2_ entered the gas channels more slowly but did reach within 8 Å of the active site in the second half of the simulations (Fig. S8b). By the end of the simulations, comparatively similar numbers of H_2_ and O_2_ molecules reached within 12-14 Å of the active site (approximately half the length of the Huc gas channel) (Table S3). However, a much smaller number of O_2_ molecules reached within 8-10 Å of the active site compared to H_2_ (Fig. S8b). This indicates that bottlenecks along the length of the tunnel selectively limit O_2_ access to the interior of Huc, which is consistent with the varying diameter of the gas channels calculated using the CAVER3 code (Fig. 3e) [56]. As discussed in the main text, an additional bottleneck in the Huc gas channel prevented O_2_ from reaching a proximity of closer than 5 Å to the Huc catalytic cluster in our simulations (Fig. S8c,d). Conversely, in a number of simulation frames H_2_ molecules are observed within 3 Å of the catalytic cluster, and in several instances in a location analogous to the ligand-bound state of the cluster (Fig. S8e,f). This indicates that the ultimate point selection against O_2_ is a bottleneck after the convergence of the three gas tunnels immediately preceding the active site entrance. Interestingly, the conformation of amino acids in this region appears responsive to whether Huc was simulated in the presence of H_2_ or O_2_. In the simulations in the presence of O_2_, arginine 443 from HucL largely adopted a conformation where it formed a close hydrogen bond with serine 104 from HucL (Fig. S8g,i). This state was never fully realized in the simulations with H_2_ and may be indicative of long-range structural changes that prime Huc for O_2_ exclusion from the catalytic site.

## Materials and Methods

### Bacterial strains and growth conditions

*M. smegmatis* mc^2^155 and its derivatives were grown on lysogeny broth (LB) agar plates supplemented with 0.05% (w/v) Tween 80 (LBT). In broth culture, *M. smegmatis* was grown in either LBT, or Hartmans de Bont minimal medium supplemented with 0.2% (w/v) glycerol, 0.05% (w/v) Tyloxapol, and 10 mm NiSO_4_ (HdB). *Escherichia coli* was maintained on LB agar plates and grown in LB broth. Selective LB or LBT medium used for cloning experiments contained gentamycin at 5 μg ml^−1^ for *M. smegmatis* and 20 μg ml^−1^ for *E. coli.* Cultures were incubated at 37 °C with rotary shaking at 150 rpm for liquid cultures unless otherwise specified. The strains of *M. smegmatis* and its derivatives and *E. coli* are listed in Table S5.

### Genetic manipulation of *M. smegmatis*

For Huc production, protein expression was performed using the *M. smegmatis* mc^2^155 strain PRC1, which lacks the glycerol response regulator *gylR (gylR* Leu154->Frameshift) and the Hhy hydrogenase *(hhyS* inactivation) [33, 75]. To facilitate Huc purification, a 2×StrepII tag was inserted at the N-terminal end of the small subunit (MSMEG_2262, HucS) through allelic exchange mutagenesis as described previously [76]. An allelic exchange construct, hucS2×StrepII (2656 bp) was synthesized by Genewiz and cloned into the SpeI site of the mycobacterial shuttle plasmid pX33 to yield the construct pHuc-2×StrepII (Table S5). pHuc-2×StrepII was propagated in *E. coli* DH5α and transformed into WT *M. smegmatis* mc^2^155 PRC1 cells by electroporation. Gentamycin was used in selective solid and liquid medium to propagate pX33. To allow permissive temperature-sensitive vector replication, transformants were incubated on LBT gentamycin plates at 28°C until colonies were visible (5–7 days). The resultant catechol-positive colonies were subcultured in LBT gentamycin broth at 40°C for 3-5 days and then diluted from broth onto fresh LBT gentamycin plates and incubated at 37 °C for 3–5 days to facilitate the integration of the recombinant plasmid, via flanking regions, into the chromosome. The second recombination event was facilitated by subculturing catechol-reactive and gentamycin-resistant colonies in LBT supplemented with 10% sucrose (w/v) for 3-5 days, followed by dilution onto LBT agar plates supplemented with 10% sucrose (w/v) and incubating at 37°C for 3–5 days. Gentamycin sensitive and catechol-unreactive colonies were subsequently screened by PCR to distinguish WT revertants from HucS-2×StrII mutants. The primers used for screening are listed in Table S5. HucM was deleted using allelic exchange mutagenesis using the same PRC1 parent strain and procedure for the 2×StrepII tagging of HucS. An allelic exchange construct consisting of ~1000bp of the genomic sequence on either side of HucM was utilized.

### Huc purification

*M. smegmatis* mc^2^155 PRC1 HucS2×StrII cells were propagated in large volumes (10-20 L) of HdB until 24 hours into stationary phase. Cells were harvested by centrifugation and resuspended in lysis buffer (50 mM Tris, 150 mM NaCl pH 8.0). Cells were lysed by cell disruptor (Emulsiflex C-5), and cell debris were removed by centrifugation at 30,000 × *g* for 15 minutes. The cell lysate was loaded onto a 1 ml StrepTrap column (Cytiva), and the column was washed extensively with lysis buffer before bound Huc was eluted with lysis buffer + 2.5 mM desthiobiotin. Huc-containing fractions from StrepTrap were determined by SDS-PAGE, pooled, and concentrated using a 100 kDa MWCO centrifugal concentrator (Amicon, Millipore). Concentrated Huc was loaded on a Superose 6 10/300 column (Cytiva), fractions containing oligomeric Huc were confirmed by SDS- and native-PAGE, pooled, and concentrated to ~6 mg ml^−1^ before flash freezing in liquid N_2_ and storage at −80°C. Typically purification yielded 10-20 μg L^−1^ of the Huc complex.

### PAGE analysis, activity staining, and western blotting

For SDS- and native-PAGE, samples were run on Bolt 4–12% SDS-polyacrylamide and native-PAGE 4-16% gels (Invitrogen) respectively, according to the manufacturer’s instructions. Gels were stained for total protein by AcquaStain Protein Gel Stain (Bulldog) and hydrogenase activity using the colorimetric electron acceptor nitrotetrazolium blue (NBT) hydrogenase. For NBT activity staining, gels were incubated in 50 mM Tris, 150 mM NaCl pH 8.0 supplemented with 500 μM NBT in an anaerobic jar amended with 7% H_2_ anaerobic mix. Incubation was conducted for 30 mins to 12 hours, depending on the level of activity. Bands exhibiting hydrogenase activity were identified based on the purple color of reduced NBT. For western blotting, proteins were transferred to nitrocellulose membrane from SDS- or native-PAGE gels using the Transblot Turbo semidry transfer apparatus (BioRad) at 25V for 30 minutes. Following transfer, the protein-containing nitrocellulose membrane was blocked with 3% (w/v) BSA in PBS (pH 7.4) with 0.1% (v/v) Tween 20 (PBST). The nitrocellulose membrane was washed three times in 20 ml of PBST and finally resuspended in 10 ml of the same buffer. Strep-Tactin HRP conjugate was then added at a 1:100,000 dilution. Bound Strep-Tactin HRP was detected by chemiluminescence using the ECL Prime detection kit (Cytiva) following the manufacturer’s specifications, blots were visualized using a ChemiDoc (BioRad).

### Amperometric measurement of Huc activity

H_2_ consumption by purified Huc was measured amperometrically with a Unisense H_2_ microsensor polarized to +100 mV for 1 hour. Initially, the electrode was calibrated with known H_2_ standards of 0%, 1%, and 10% H_2_ (v/v) in gas saturated buffer (50 mM Tris, 150 mM NaCl, pH 8.0). This buffer was prepared by firstly bubbling all buffers with 100% nitrogen gas for 1 hour to remove any traces of oxygen, then bubbling 100% H_2_ (v/v) through the de-oxygenated buffer for 10 minutes. Prior to degassing and regassing, all buffers contained electron acceptors to a final concentration of 200 μM (menadione, NBT, or benzyl viologen). For each reading, 1 ml of 1% (8 μM) H_2_ infused buffer containing the electron acceptor was added to the microrespiration assay chamber, to a final concentration of 1% H_2_. The electrode was subsequently placed into the chamber to equilibrate. Once equilibrated (approximately 10 minutes), BSA (0.3 mg ml^−1^) and Huc (1-3 nM in 0.3 mg ml^−1^ BSA) were added to the chamber via a needle so as not to disrupt the gas balance in the solution. The changes in H_2_ concentration were measured using the Unisense Logger Software and linear rates of H_2_ consumption were measured from the addition of Huc until complete H_2_ consumption. The linear rate of H_2_ consumption by Huc was calculated between a concentration of approximately 2.5 μM H_2_ to 0.0125 μM H_2_ over 8-time points. These rates of H_2_ consumption were used to plot a Michaelis-Menton curve and calculate Huc-specific activity in the presence of different electron acceptors, as well as the maximal velocity of Huc.

### Mass spectrometric menaquinone detection

Samples were prepared for liquid chromatography-mass spectrometry (LC-MS) analysis using a modified Folch extraction [77]. Briefly, a 100 μL solution of purified protein (~57 μg) was treated with 2000 μL of 2:1 chloroform:methanol v/v after which the mixture was shaken for 10 minutes and allowed to stand for a further 50 minutes. 400 μL of water was added and the mixture was shaken for 10 minutes after which the sample was allowed to stand until the two phases had completely separated. The lower chloroform-rich phase was then transferred to a 2 mL sample vial and the solvent was removed under a stream of nitrogen gas. The resulting residue was reconstituted in 100 μL of 2:1 chloroform:methanol v/v and transferred to a 200 μL sample insert, the solvent was again removed and the sample reconstituted in a 7:3 mixture of LC solvent A:LC solvent B v/v. Samples were analysed using a Dionex RSLC3000 UHPLC (Thermo) coupled to a Q-Exactive Plus Orbitrap MS (Thermo) using a C18 column (Zorbax Eclipse Plus C18 Rapid Resolution HD 2.1 x 100mm 1.8 micron, Agilent) with a binary solvent system; solvent A = 40% isopropanol and solvent B = 98% isopropanol both containing 2mM formic acid and 8 mM ammonium formate. Linear gradient time-%B as follows: 0 min-0%, 8 min-35%, 16 min-50%, 19 min-80%, 23 min-100%, 28 min-100%, 30 min-0%, 32 min 0%. The flow rate was 250 μL min^−1^, the column temperature 50°C, and the sample injection volume was 10 μL. The mass spectrometer operated at a resolution of 70,000 in polarity switching mode with the following electrospray ionization source conditions: Spray voltage 3.5kV; capillary temperature 300 °C; sheath gas 34; Aux gas 13; sweep gas 1 and probe temp 120 °C.

### Mass spectrometric identification of HucM

The 18 kDa band on SDS-PAGE originating from the Huc complex was excised and extracted from the gel matrix. Upon destaining, the proteins were reduced with tris(2-carboxyethyl)phosphine (Pierce), alkylated with iodoacetamide (Sigma), and digested with mass spectrometry grade trypsin (Promega). The extracted peptides were analyzed by LC-MS/MS on an Ultimate 3000 RSLCnano System (Dionex) coupled to an Orbitrap Fusion Tribrid (ThermoFisher Scientific) mass spectrometer equipped with a nanospray source. The peptides were first loaded and desalted on an Acclaim PepMap trap column (0.1 mm id × 20 mm, 5 μm) and then separated on an Acclaim PepMap analytical column (75 μm id × 50 cm, 2 μm) over a 30 min linear gradient of 4–36% acetonitrile/0.1% formic acid. The Orbitrap Fusion Tribrid was operated in data-dependent acquisition mode with a fixed cycle time of 2 s. The Orbitrap full ms1 scan was set to survey a mass range of 375–1800 m/z with a resolution of 120,000 at m/z 400, an AGC target of 1 × 10^6,^ and a maximum injection time of 110 ms. Individual precursor ions were selected for HCD fragmentation (collision energy 32%) and subsequent fragment ms2 scan were acquired in the Orbitrap using a resolution of 60,000 at m/z 400, an AGC target of 5 × 10^5,^ and a maximum injection time of 118 ms. The dynamic exclusion was set at ±10 ppm for 10 s after one occurrence. Raw data were processed using Byonic (ProteinMetrics) against a protein database covering *Mycobacterium smegmatis* mc^2^155. The precursor mass tolerance was set at 20 ppm, and fragment ions were detected at 0.6 Da. Oxidation (M) was set as dynamic modification, carbamidomethyl (C) as fixed modification. Only peptides and proteins falling below a false discovery rate of 0.01 were reported.

### Gas chromatography analysis

Gas chromatography experiments were used to measure the consumption of H_2_ gas by both pure Huc protein and cultures of *M. smegmatis* expressing Huc. For pure protein experiments, 5 ml of buffer (50 mM Tris, 150 mM NaCl, pH 8.0, 0.3 mg ml^−1^ BSA) with 200 μM NBT was contained in a 120 ml sealed serum vial, in triplicate. The headspace of the vial was flushed with 10 ppm H_2_, in an atmospheric gas mix, for 10 mins and then 50 nM Huc was added via a syringe to begin the reaction. Vials were incubated at room temperature, stirring, and hydrogen gas concentration was monitored using a pulsed discharge helium ionisation detector (model TGA-6791-W-4U-2, Valco Instruments Company Inc.) over specific time intervals for approximately 30 hours. For pure culture experiments 30 ml of cells in HdB minimal media, grown to one-day post-OD maximum, were flushed with 10 ppm H_2_ in 120 ml sealed serum vials and incubated shaking at 37°C and 150 rpm. H_2_ readings were taken as described for pure protein. Controls of buffer with no Huc for the pure protein experiments and heat-killed (120°C, 20 mins) culture for the pure culture experiments were used. All curves were calibrated with standards of N_2_ (0 ppm H_2_), pure air (0.53 ppm H_2_), and 10 ppm H_2_. Data were plotted as time versus hydrogen concentration.

### Differential scanning fluorimetry

To determine the stability of the Huc complex, thermal melting was performed from 20 to 90°C using a PrometheusNT.48 DSF (Nanotemper) using high sensitivity capillaries. The ratio of change in fluorescence at 330 and 350nm was monitored to determine protein unfolding. Melting was performed with a Huc concentration of 0.2 mg ml^−1^ in buffer containing 50 mM Tris, 150 mM NaCl pH 8.0.

### CryoEM imaging

Samples (3 μl) were applied onto a glow-discharged UltrAuFoil grid (Quantifoil GmbH) and were flash-frozen in liquid ethane using the Vitrobot mark IV (Thermo Fisher Scientific) set at 100% humidity and 4 °C for the prep chamber. Data were collected on a Titan Krios microscope (Thermo Fisher Scientific) operated at an accelerating voltage of 300 kV with a 50 μm C2 aperture at an indicated magnification of 105 K in nanoprobe EFTEM mode. Gatan K3 direct electron detector positioned post a Gatan BioQuantum energy filter, operated in a zero-energy-loss mode with a slit width of 25 eV was used to acquire dose fractionated images of the Huc complex without an objective aperture. Two initial Huc datasets were collected, one low concentration dataset (0.4 mg ml^−1^) composed of 2,226 movies, and a medium concentration dataset (1 mg ml^−1^) composed of 3,113 movies. Movies were recorded in hardware-binned mode (previously called counted mode on the K2 camera) yielding a physical pixel size of 0.82 Å pixel^−1^ with an exposure time of 6 s amounting to a total dose of 66.0 e^−^ Å^−2^ at a dose rate of 1.5 e^−^ pixel^−1^ s^−1^, which was fractionated into 60 subframes. A further high concentration dataset (4 mg ml^−1^) of 9868 micrographs was also recorded using the same microscope but in ‘super-resolution’ mode on the K3 detector, the physical pixel size was 0.42 Å with an exposure time of 4 s amounting to a total dose of 60.4 e^−^ Å^−2^ at a dose rate of 1.59 e^−^ pixel^−1^ s^−1^, which was fractionated into 60 subframes. Defocus range was set between −1.5 and −0.5 μm.

### CryoEM data processing and analysis

Micrographs from all datasets were motion-corrected using UCSF Motioncor and dose weighted averages had their CTF parameters estimated using CTFFIND 4.1.8, implemented using Relion 3.1.2[78].

#### Huc oligomer reconstruction from Dataset1

Particle coordinates were determined by crYOLO 1.7.6 using a model trained on manually particles picked from 20 micrographs [79]. Unbinned particles were extracted from micrographs using Relion 3.1.2, before being imported into cryoSPARC 3.3.1 for initial 2D classification to remove bad particles, followed by ab initio model generation and 3D refinement [80]. Refined particles were reimported into Relion 3.1.2 and CTF refinement was performed, followed by Bayesian Polishing [78]. Particles were reimported into cryoSPARC 3.3.1 for final 2D classification to remove residual bad particles, followed by non-uniform 3D consensus refinement to generate final maps at 2.19 Å (FSC = 0.143, gold standard).

#### Huc2S2L subunit Dataset 1

HucS_2_L_2_ particles were picked using gautomatch V 0.53 (developed by Kai Zhang, MRC Laboratory of Molecular Biology, Cambridge, UK) using a diameter of 80 Å and a minimum distance between picked particles set to 20 Å. The resultant particles were extracted and binned 4 times to a box size of 56 pixels. A small subset of unbinned particles with 224 pixel box size were subjected to 2D classification to exclude noisy particles and the remainder were selected and subjected to 3D ab initio model generation with 4 classes using cryoSPARC v3.0.1 [80]. A class corresponding to a population of Huc2S2L subunits showed clear C2 symmetry and this particle set was subjected to homogenous refinement with C2 symmetry applied resulting in a 3.25 Å resolution reconstruction. The resulting volume was used as the initial model for further classification and refinement of the full dataset as shown in the workflow Fig. S5. Particles corresponding to the subunits were selected from binned dataset following 2D classification and heterogenous refinement. The cryoSPARC 3D alignments from heterogenous refinement were then exported to Relion using the csparc2star.py script in the pyem v0.5 package [81]. In Relion, particles were re-extracted with no binning using the refined coordinates from the imported particles. These were re-imported into cryoSPARC and subjected to heterogeneous refinement followed by homogenous refinement with C2 symmetry applied. The FSC from the resultant reconstruction showed particle duplication as a result of over-picking using gautomatch. The refined particle coordinates were imported into Relion as described above and were cleaned to remove duplicated particles. The cleaned re-extracted particles were subjected to a round of CTF refinement followed by auto-refinement and Bayesian polishing. The resultant polished particles were imported back into cryoSPARC and CTF refinement, and non-uniform refinement with CTF parameter optimization was performed to reach a final global resolution of 1.67 Å (FSC = 0.143, gold standard), which was close to the Nyquist limit for the dataset (1.64 Å).

#### Huc2S2L subunit Dataset2

A second dataset at higher magnification was collected to overcome the Nyquist limit imposed on the map resolution of Dataset 1. The particles were picked, extracted, and binned as described above and in Fig. S4. Multiple rounds of 2D classification, ab initio classification, and heterogeneous refinement was performed to retain only particles containing the full Huc oligomer. Homogenous refinement was performed on the binned particles with the Huc2S2L subunit map as an initial model in order to centre the particles on the Huc2S2L subunit. These particles were then re-extracted in Relion based on the refined coordinates to a box size of 380 pixels. Homogenous refinement with C2 symmetry resulted in a subunit reconstruction map of 2.15 Å. This step ensured the retention of all *Huc2S2L* particles with a good signal to noise ratio. In order to ensure that all symmetry-related *Huc2S2L* particles corresponding to the Huc oligomer are retained, the above-selected particles were reextracted with a larger box size of 512 pixels and rescaled to 128 pixels (4x binned). The resultant particle set was subjected ab initio model generation followed by homogenous refinement using C4 symmetry to yield a 4.10 Å map of the Huc oligomer. The particles were re-extracted based on the refined coordinates from the previous step at a box size of 576 pixels and then rescaled to 288 pixels (2x binned). These particles were then subjected to homogenous refinement followed by CTF refinement and Non-Uniform refinement in cryoSPARC to yield a 2.05 Å resolution map [80]. The refined particle set was then re-imported into Relion, and Bayesian polishing was performed. The resultant ‘shiny’ particles were extracted to a final box size of 576 pixels with no binning and were re-imported into cryoSPARC to perform symmetry expansion and final refinement. Duplicate particles were excluded to retain 153,359 Huc oligomers. Resultant particles were subjected to one round of refinement imposing C4 symmetry, which yielded a 2.19 Å map. The resultant refined particles were symmetry expanded and subjected to local refinement within cryoSPARC 3.3.2 with no symmetry to yield a 2.16 Å map for *Huc2S2L.* The resultant vol and particles were aligned to C2 symmetry axis to exploit the C2 symmetry within the *Huc2S2L.* The symmetry aligned particles were then refined with C2 symmetry followed by iterative Global CTF refinement, per particle defocus refinement followed by Ewald Sphere correction and a final round of per particle defocus refinement to reach a final global resolution of 1.52 Å (FSC = 0.143, gold standard).

#### HucM tail reconstruction

The polished Huc oligomer particles from dataset2 were subjected to ab initio model generation with 3 classes to retain only the particles that showed clear density for the HucM tail region. The selected particles were then ctf refined and Non-Uniform refined to yield 2.09 Å map. The stalk density was isolated using volume tools in Chimera and low pass filtered to 20 Å to derive the initial model for local refinement in cryosparc [82]. A dilated cosine padded soft mask was generated using the initial model. The coordinates at the attachment of the stalk to the main body of the subunit were determined using the volume tracer tool in Chimera and used to define new fulcrum point for localised refinement (Fig. S6). Two rounds of masked local refinement resulted in improving the density of the stalk region associated to membrane as shown in the figure to a resolution of 5.7 Å (no mask). The resultant FSC calculates a map resolution of 5.21 Å (FSC = 0.143, gold standard) but the map features are more consistent with a resolution of 6-8 Å on visual inspection.

#### HucM tail reconstruction

The particle set resulting in a 2.15 Å map during the Huc2S2L subunit reconstruction from Dataset 2 was re-extracted at a box size of 512 pixels and rescaled to 128 pixels (4x binned). The resultant particle set was subjected to 2D classification followed by ab initio model generation and homogenous refinement using C4 symmetry to yield a 4.10 Å map of Huc oligomer. The particles were re-extracted based on the refined coordinates from the previous step at a boxsize of 576 pixels and then rescaled to 288 pixels (2× binned). These particles were then subjected to homogenous refinement followed by CTF refinement and Non-Uniform refinement in cryoSPARC to yield 2.05 Å resolution map [80]. The refined particles set was then re-imported into Relion and Bayesian polishing was performed, the resultant ‘shiny’ particles were extracted to a final box size of 576 pixels with no binning. These particles were subjected to ab initio model generation with 3 classes to retain particles that showed clear density for the HucM tail region. The selected particles were then ctf refined and Non-Uniform refined to yield 2.09 Å map. The stalk density was isolated using volume tools in Chimera and low pass filtered to 20 Å to derive the initial model for local refinement in cryosparc [82]. A dilated cosine padded soft mask was generated using the initial model. The coordinates at the attachment of the stalk to the main body of the subunit were determined using the volume tracer tool in Chimera and used to define new fulcrum point for localised refinement (Fig. S6). Two rounds of masked local refinement resulted in improving the density of the stalk region associated to membrane as shown in the figure to a resolution of 5.7 Å (no mask). The resultant FSC calculates a map resolution of 5.21 Å (FSC = 0.143, gold standard) but the map features are more consistent with a resolution of 6-8 Å on visual inspection.

### Huc model building and visualization

A model of the HucS and HucL subunit dimers was generated using Alphafold and docked into one-half of the high-resolution Huc Dimer maps using ChimeraX [49, 83]. The model was refined and rebuilt into map density using Coot [84]. [NiFe], [3Fe-4S], and menaquinone cofactors associated with Huc were downloaded from the PDB and customized restraints generated using Elbow within the PHENIX package before they were fitted and refined into maps using Coot [85, 86]. The model was then refined using real-space refinement within PHENIX [87]. Once model building was complete the model was symmetry expanded using the Map symmetry tool and waters were added using DOUSE within the PHENIX package [86]. The refined dimer model was docked into the Huc oligomer map using ChimeraX [83], with HucM manually built into the available density for one subunit using Coot [84], followed by iterative real-space refinement within PHENIX and further model building. Once model building was complete the model was symmetry expanded using the Map symmetry tool within the PHENIX package [86]. Model quality was validated using MolProbity [88]. Images and movies were generated in Coot, ChimeraX, and Pymol [83, 84].

### AlphaFold structural modeling

AlphaFold modeling was performed using AlphaFold version 2.1.1 implemented on the MASSIVE M3 computing cluster [49]. The sequence of the HucM C-terminal region (amino acids 80-189) was provided and modeling was run in multimer mode, with four molecules of HucM requested. The five ranked models produced by AlphaFold were compared for consistency with the top-ranked model utilized for further analysis and figure generation.

### Molecular dynamics parameterization and simulation

The 1.52 Å crystallographic structure of Huc hydrogenase was used to construct the simulation system using the Charmm36m forcefield [89, 90] with the TIP3P [91] water model. Sodium counterions were added to the solution to neutralize the system charge. Three sets of simulations were set up containing no gas molecules or containing 250 hydrogen or 250 oxygen molecules enriched in solution. Gas molecules were randomly inserted into the water phase. Each simulation was initiated with different starting velocities and gas positions and 3 independent simulations of each system were performed. Simulations were run for 50 ns. Molecular visualization and analyses were performed using VMD software [92] The topologies of the metal cofactors were constructed based on the coordinates from the CryoEM structure. Previous work on hydrogenase cofactors [93] has demonstrated the transferability of structurally identical metallic cofactors between hydrogenases. Thus, atomic partial charges and force constants for the bond lengths of the metal cofactors were taken from previous studies of the structurally similar *D. fructosivarans* [NiFe] hydrogenase [93]. Equilibrium values of bond lengths and angles within the metal cofactor were taken directly from the CryoEM structure. MD simulations were performed in a single redox state. The ready active NI-S form of the active site and oxidized form of the iron-sulfur clusters were chosen based on previous similar analyses [94]. The [NiFe] cofactor had bond constraints placed across the CN and CO ligands as well as between the iron atom and two bound sulfur atoms. Lennard-Jones parameters were taken from the Charmm36m [89, 90] forcefield where appropriate, with exception of the cyanuric carbon [95] with the iron parameters taken from physical data to have a non-zero epsilon value [96]. The QL molecular oxygen model developed by Javanainen et al [97] was chosen to better define the quadrupole within the molecule. The oxygen atoms are given mass and non-zero Lennard-Jones parameters as well as a partial charge. A massless virtual site, OD1, exists at the midpoint of two oxygen atoms to carry the neutralizing charge, simulating the charge distribution within the oxygen quadrupole. Likewise, the hydrogen model was given mass and charge distribution coherent with the work of Hunter et al. [94] MD simulations were performed via the GROMACS v 2021.3 simulations suite [98]. Energy minimization was performed using the steepest descent followed by a second minimization using the conjugant gradient approach, both set to a maximum force of 1000 kJ mol^−1^ nm^−1^. Equilibration simulations were performed within the NVT ensemble at 310 K. Position restraints were applied to the protein, cofactors and magnesium ion, with a 10^5^ kJ/(mol nm^2^) force constant for 100 ps. Further equilibration was carried out within the NPT ensemble with pressure maintained at 1 bar using the Parrinello-Rahman barostat, [99] for 100 ps. Production simulations in the NPT ensemble were each of duration 50 ns, at a temperature of 310 K. Long-range electrostatics were treated with the Particle Mesh Ewald method [100], with values for the real space long range cut-off for these and Van der Waals forces being set to 10 Å. Covalent bonds containing hydrogen atoms were restrained using the LINCS algorithm [101, 102] allowing a 2 fs integration step.

### Fourier transform infrared spectroscopy

4 μl of 5 mg ml^−1^ Huc enzyme solution in a buffer containing 50 mM Tris, 150 mM NaCl pH 8 was deposited on an ATR crystal surface. The sample was applied under laboratory atmosphere (air), dried under 100% nitrogen gas, and rehydrated with a humidified aerosol (100 mM Tris-HCl (pH 8)) as described previously [103, 104]. A custom build PEEK cell (inspired by Stripp et al. [105]) that allows for gas exchange sealed the ATR unit (BioRadII from Harrick) mounted in the FTIR spectrometer (Vertex V70v, Bruker). Spectra were recorded with 2 cm^−1^ resolution, 80 Hz scanner velocity, and averaged over a varying number of scans (at least 100 Scans). All experiments were performed at ambient conditions (room temperature and pressure, hydrated enzyme films). Data were analyzed using Qsoas and Origin 8 software. To assess the response of the Huc [NiFe] cluster to ambient air, N_2_, and H_2_ the PEEK cell was flushed sequentially with each gas in the following sequence: N_2_ (ambient air isolated enzyme), ^2^H_2_, N_2_, H_2_, ambient air. Changes in the Huc spectra were allowed to stabilize before the spectra in each gas were collected.

### Protein film electrochemistry

Protein film electrochemistry experiments were carried out under anaerobic conditions. The three-electrode system was made up of (1) Ag/AgCl (4 M KCl) as reference electrode, (2) rotating disk 5 mm OD pyrolytic graphite edge (PGE) plane (epoxy encapsulated) as working electrode, and (3) graphite rod as the counter electrode. The glass cell used featured a water jacket for temperature control and a cell gas inlet/outlet for H_2_ flow. The buffer used was composed of 5 mM MES, 5 mM CHES, 5 mM HEPES, 5 mM TAPS, 5 mM sodium acetate, with 0.1 M Na_2_SO_4_ as carrying electrolyte titrated with H_2_SO_4_ to pH 7.0, and purged with N_2_ for 3 to 4 hours. To remove residual O_2_ in the PGE electrode, cyclic voltammograms were run at 100 mV/s from 0 to −800 mV (vs. NHE) for 10 scans. To adhere Huc to the deaerated PGE electrode the surface was abraded with P1200 sandpaper and rinsed with purified water. 5 mg ml^−1^ Huc enzyme solution in a buffer containing 50 mM Tris, 150 mM NaCl pH 8 was mixed with polymyxin B sulfate and transferred to the electrode surface. The cell solution was then saturated with H_2_ (1 atm, 10 min) before the cyclic voltammogram of the system with the immobilized enzyme was recorded at 10 mV/s. The cyclic voltammogram of the blank electrode (no enzyme immobilized) was then recorded at 10 mV/s. Electrochemical data were acquired using an Eco/Chemie PGSTAT10 and the GPES software (Metrohm/Autolab). Data were analyzed using Qsoas and Origin 8 software. All potential values are referenced versus NHE. Experiments were conducted on two independent Huc films, with each film scanned three times.

